# The PP2A/4/6 subfamily of phosphoprotein phosphatases regulates DAF-16 during aging and confers resistance to environmental stress in postreproductive adult *C. elegans*

**DOI:** 10.1101/2020.02.18.953687

**Authors:** Rebecca S. Rivard, Julia M. Morris, Matthew J. Youngman

## Abstract

Insulin and insulin-like growth factors are longevity determinants that negatively regulate Forkhead box class O (FoxO) transcription factors. In *C. elegans* mutations that constitutively activate DAF-16, the ortholog of mammalian FoxO3a, extend lifespan by two-fold. While environmental insults induce DAF-16 activity in younger animals, it also becomes activated in an age-dependent manner in the absence of stress, modulating gene expression well into late adulthood. The mechanism by which DAF-16 activity is regulated during aging has not been defined. Since phosphorylation of DAF-16 generally leads to its inhibition, we asked whether phosphatases might be necessary for its increased transcriptional activity in adult *C. elegans*. We focused on the PP2A/4/6 subfamily of phosphoprotein phosphatases, members of which had been implicated to regulate DAF-16 under low insulin signaling conditions but had not been investigated during aging in wildtype animals. Using reverse genetics, we functionally characterized all *C. elegans* orthologs of human catalytic, regulatory, and scaffolding subunits of PP2A/4/6 holoenzymes in postreproductive adults. We found that PP2A complex constituents PAA-1 and PPTR-1 regulate DAF-16 during aging and that they cooperate with the catalytic subunit LET-92 to protect adult animals from ultraviolet radiation. PP4 complex members PPH-4.1/4.2, SMK-1, and PPFR-2 also appear to regulate DAF-16 in an age-dependent manner, and they contribute to innate immunity. Interestingly, SUR-6 but no other subunit of the PP2A complex was necessary for the survival of pathogen-infected animals, suggesting that a heterotypic PP4 complex functions during aging. Finally, we found that PP6 complex constituents PPH-6 and SAPS-1 contribute to host defense during aging, apparently without affecting DAF-16 transcriptional activity. Our studies indicate that a set of PP2A/4/6 complexes protect adult *C. elegans* from environmental stress, thus preserving healthspan. Therefore, along with their functions in cell division and development, the PP2A/4/6 phosphatases also appear to play critical roles later in life.

## Introduction

Perhaps more than any other, the Insulin and IGF-signaling (IIS) pathway has been unequivocally established as a primary modulator of lifespan across evolutionarily diverse species (1). Genetic and behavioral changes that modify IIS pathway activity change the quality and length of life dramatically. Loss-of-function mutations in the insulin receptor correlate with a 2- to 3-fold extension in the lifespan in some species (2). Moreover, insulin signaling mutants appear more youthful and are more resistant to a variety of environmental insults, demonstrating a connection between stress resistance and lifespan (3, 4). Thus, the ability of an organism to withstand encounters with stressful stimuli or to otherwise neutralize the consequent damage associated with acute stress is a primary determinant of its overall health during aging and, ultimately, how long it lives. The IIS pathway, and its ultimate target, the Forkhead box family (FoxO) transcription factor, have a profound influence on lifespan and stress resistance. Consequently, understanding their regulation over time could suggest the basis for the milestones of aging and reveal potential avenues for manipulating the rate of aging.

In *C. elegans* the FoxO protein DAF-16 is a transcriptional regulator that determines lifespan in part by conferring resistance to environmental insults that have the potential to cause catastrophic cellular and molecular damage. As the ultimate downstream target of the insulin and IIS pathway, the transcriptional activity of DAF-16 is inhibited when insulin-like ligands are bound to the insulin receptor, DAF-2. In the absence of these soluble ligands, DAF-16 is free to translocate to the nucleus where it upregulates the expression of genes that promote longevity, and it is because of this function of DAF-16 that *daf-2* mutants live twice as long as their wildtype counterparts (5). DAF-16 activity is also triggered even if the IIS pathway is intact when animals encounter harmful stimuli including high temperature and ultraviolet radiation (6). Under these stress-inducing conditions, DAF-16 targets may include genes that encode detoxifying enzymes, immune effectors, and chaperones that restore homeostasis, preserve cellular health and contribute to host defense (7). Recently reported evidence suggests that DAF-16 may be activated in an age-dependent manner, despite the absence of acute stress. During adulthood, the expression levels of the *daf-16a* isoform, and to an even greater extent the *daf-16d* isoform progressively increase (8). This is coupled to increased expression of a subset of DAF-16 transcriptional targets in adults as compared to in larvae, suggesting that DAF-16 is activated as *C. elegans* age (9). How DAF-16 activity might be regulated in an age-dependent manner and whether that regulatory mechanism is unique to aging is not known.

Studies of larval stage worms and especially of *daf-2* mutants have revealed that the transcriptional activity of DAF-16 is regulated on two primary levels. First, inside the nucleus DAF-16 associates with other proteins that modulate its transcriptional output. For example, HCF-1 binds to DAF-16 and interferes with its ability to bind to the promoters of certain target genes (10). Conversely, the basic helix-loop-helix transcription factor HLH-30 forms a complex with DAF-16, and the two proteins together then co-regulate a subset of targets enriched for genes that promote longevity. The very presence of DAF-16 inside the nucleus is subject to control from multiple inputs, representing a second level of regulation. The nuclear localization of DAF-16 is inhibited predominantly through the IIS pathway. Specifically, IIS signaling activates the evolutionarily conserved kinase AKT-1 which phosphorylates DAF-16. Assuming that this modification of DAF-16 has the same effect that it does when FoxO transcription factors in other species are phosphorylated, a binding site for 14-3-3 proteins is created and the nuclear export signal of DAF-16 is exposed, leading to its expulsion from the nucleus and sequestration in the cytosol (11). Supporting this possibility, DAF-16 accumulates in the nuclei of larval worms that lack the 14-3-3 proteins PAR-5 and FTT-2 (12, 13). The inhibitory phosphorylation imparted to FoxO transcription factors by AKT is opposed by the action of phosphoprotein phosphatase PP2A. In mammals, both FoxO1 and FoxO3a are substrates of PP2A, and biochemical evidence indicates that PP2A selectively dephosphorylates the AKT phosphorylation sites of FoxO family members (14–17). Moreover, PP2A also dephosphorylates AKT itself, thereby inactivating it (18–20). This particular mechanism of PP2A functioning as an indirect positive regulator of FoxO transcription factors appears to be conserved in *C. elegans*. In *daf-2* mutants, the constitutive activation of DAF-16 requires PPTR-1, a conserved regulatory subunit of PP2A complexes (21). PPTR-1 associates with AKT-1 but not DAF-16 in worms, and overexpression of PPTR-1 reduces the levels of phosho-AKT-1. Since as a whole the expression levels of insulin-like peptides considered to be agonists of DAF-2 either remain constant or increase during adulthood, the presence of a sustained inhibitory signal through the IIS pathway that persists in adult animals seems likely (9). A regulator that can counteract this signal is therefore an attractive candidate for functioning to modulate the activity of DAF-16 during aging, and this is what led us to develop the hypothesis that phosphoprotein phosphatases modulate DAF-16 transcriptional activity in an age-dependent manner.

In mammals, PP2A may generally be considered to play an anti-aging role, in part because it protects organisms from age-related disease. For example, PP2A is the primary phosphatase that dephosphorylates Tau protein in the brain and therefore helps to prevent the neurodegeneration that leads to Alzheimer’s disease (22). The level and activity of PP2A detected in postmortem brains of Alzheimer’s patients are, in fact, reduced compared to healthy controls (23). Similarly, low PP2A activity in monkey brains is associated with elevated levels of phosphorylated alpha-synuclein, a hallmark of Parkinson’s disease (24, 25). Outside of the context of acute disease, PP2A activity in the liver of SAMP8 mice is reduced and is correlated with increased levels of inactive, phosphorylated FoxO1 which may be responsible for the animals’ shortened lifespan (26).

PP2A is part of a subfamily of phosphoprotein phosphatases that also includes PP4 and PP6. Phosphoprotein phosphatases (PPPs) comprise a large class of evolutionarily conserved enzymes, that antagonize kinases by removing phosphate groups from a broad repertoire of substrates. The PPP class enzymes are structurally distinct from the PPM Ser/Thr phosphatases in that members of the PPM family are Mg^2+^-dependent, unlike members of the PPP family(27). PPPs can be divided into two evolutionarily distinct branches composed of PP1/PP2A/PP2B and PP5/PPEF/PP7 based on divergent regulatory and catalytic domains, with the PP2A subgroup containing the PP2A/PP4/PP6 subfamily(27). PPP family phosphatase active sites are similar, but differ in sensitivity and response to certain small molecule inhibitors(28). While PP3/5/7 contain N- or C-terminal domains that act as built in regulatory factors, the PP1/2A/4/6 catalytic subunits all have a highly conserved globular domain that acts in concert with one (PP1/6) or two (PP2A/4/6) binding partners. Catalytic subunits of the PP2A/4/6 subfamily associate with regulatory proteins and, in some cases, a scaffolding protein to comprise a functional holocomplex. Multiple versions of the PP2A/4/6 complexes are possible because there are several genes whose products bind and regulate the same catalytic subunit(29). For example, in humans there are 18 different regulatory proteins capable of associating with the PP2A enzyme, allowing for the generation of nearly 100 holoenzymes (30, 31). The combinatorial diversity of these complexes is believed to contribute to the subcellular localization, substrate specificity, and thus the functional capacity of the phosphoprotein phosphatases, depending on their subunit composition(32).

In evolutionarily diverse species, the PP2A/4/6 subfamily regulates a wide array of processes, including cell cycle progression, meiosis, cellular differentiation, and the activity of multiple signaling pathways including the IIS pathway (30,33–42). Interestingly, in *C. elegans* PP2A is not the only member of the PP2A/4/6 family with a connection to the insulin signaling pathway. Just as the PP2A subunit PPTR-1 is required for the extended lifespan of *daf-2* mutants, so too is the regulatory subunit of the PP4 complex SMK-1 (43). SMK-1 is orthologous to human PP4R3, *D. melanogaster* flfl, and *S. cerevisiae* Psy2p. During embryonic development in worms, SMK-1 associates with replicating chromatin, triggering the recruitment PPH-4, the catalytic subunit of the PP4 complex, and silencing the CHK-1-mediated response to DNA damage, allowing cell cycle progression(44). Outside of the context of embryonic development, SMK-1 appears to be constitutively expressed in the nucleus of several tissues where it plays a role in longevity and stress resistance. Reminiscent of phenotypes associated with inhibiting PPTR-1, when *smk-1* expression is blocked by RNAi, *daf-2* animals are no longer resistant to a variety of stresses (43). Furthermore, the expression of a subset of DAF-16 transcriptional targets is reduced following RNAi against *smk-1* because of a deficiency in transcription initiation (45). This apparent functional overlap between PP2A and PP4 in regulating DAF-16 in a genetic background where DAF-16 is hyper-activated raises the question as to which member is principally responsible for activating DAF-16 during the aging process in wildtype animals. Moreover, considering that different combinations of subunits allow for the possibility of multiple versions of the PP2A/4/6 complexes, a more nuanced version of this question would ask about the specific identity of the individual constituents of the complex(es) responsible for regulating DAF-16 during adulthood. This is what our study sought to address.

Here we present the first systematic investigation of the role of the PP2A/4/6 subfamily of phosphoprotein phosphatases during aging in wildtype *C. elegans*. We used a reverse genetic approach to functionally characterize all *C. elegans* orthologs of human PP2A/4/6 catalytic, scaffold, and regulatory subunits in juvenile and postreproductive adult animals. Our data suggest that the PP2A and PP4 complexes regulate the age-dependent activity of DAF-16. Moreover, we present evidence to support the existence of multiple versions of PP2A, PP4, and PP6 complexes with apparent specialized functions to protect adult *C. elegans* from different environmental stresses and thus as a group contribute to preserving healthspan.

## Methods

### C. elegans *growth and maintenance*

Bristol N2 and VIL001 *mjyIs001* [*Plys7::gfp*] *C. elegans* strains were grown and maintained under standard laboratory conditions (46). To generate strain VIL001, stable transgenic animals harboring an extrachromosomal *lys-7* promoter::gfp fusion construct (47) were subjected to gamma irradiation to yield a chromosomal integration of *Plys7::gfp*. F_2_ segregants of the irradiated P_0_ animals yielding 100% GFP-expressing progeny were selected for further analysis. One of these lines was backcrossed to the N2 wildtype strain seven times and then designated VIL001.

### *C. elegans* synchronization

Worms were synchronized via sodium hypochlorite treatment (48). After hatching overnight in sterile M9 buffer, approximately 2000 L1 stage worms were dropped to fresh, pre-seeded RNAi plates or to NGM plates seeded with *E. coli* OP50 (see below).

#### RNAi treatment

Animals were treated with RNAi through their food source as described by Fraser *et al.* with modifications as described below (49). All RNAi sequences were confirmed by sequencing prior to use. Bacteria from the Source BioScience library were streaked onto LB plates containing ampicillin (amp, 5µg/mL) and tetracycline (tet, 1.25µg/mL) and allowed to grow overnight at 37°C. One colony from each plate was used to inoculate 200mL of LB with amp (1ng/mL) and grown overnight in a 37°C shaker. Bacteria were centrifuged at 5,000g for 10 minutes, resuspended in 20mL of LB containing amp, and 1mL was dropped onto individual RNAi plates (standard NGM plates with 1mg/mL carbenicillin, 2µM IPTG). Lawns of *E. coli* RNAi clones were allowed to grow for at least two days before worms were introduced to the plates. Except in the cases where either *let-92* or *paa-1* were the targets of the treatment, synchronized cohorts of L1 larvae were dropped onto seeded RNAi plates and maintained at 20°C. When these animals reached the L4 stage they were transferred to pre-seeded RNAi plates containing 25 µg/mL FUdR (5-fluorodeoxyuridine). Since initiating knockdown of *let-92* and *paa-1* at L1 prevented maturation past L4, RNAi treatment targeting these genes was delayed until animals had reached L4 on NGM plates seeded with OP50. Worms subjected to heat stress or UV irradiation remained on RNAi plates until they died. For bacterial infection assays, worms were transferred from the RNAi plates at either the L4 stage or at Day 6 of adulthood to plates containing *Pseudomonas aeruginosa* where they remained throughout the duration of the assay.

### Pseudomonas aeruginosa (PA14) *infection assays*

Infection assays were carried out as described in Tan *et al.* with the following modifications (50). For each infection assay replicate worms were cultured on RNAi plates as described above and approximately 30 animals were transferred to each of three slow kill assay 3.5 cm plates at L4 and Day 6 where they were maintained at 25°C for the duration of the experiment. These plates contained 25 µg/mL FUdR, were seeded with 6 µl of an overnight culture of PA14, and then incubated for one night at 37°C and a second night at room temperature prior to the start of the infection assay. Worm survival was scored in the morning and evening every day by counting the number of animals that responded to gentle prodding with a wire pick. Worms that failed to move were scored as dead and removed from the plates.

#### Thermotolerance assay

After being synchronized by sodium hypochlorite treatment as described above, L1 larvae were dropped on to RNAi plates seeded with a clone expressing dsRNA to target a given gene of interest. Worms were maintained on those plates until the L4 larval stage when they were split into two groups. The first group of worms was subjected to thermal stress as described previously with some modifications (43). Briefly, approximately 100 L4 worms were distributed evenly between three pre-seeded 3.5 cm RNAi food plates containing FUdR and then incubated at 35°C for the duration of the heat treatment and scored every few hours for survival with animals being counted as alive if they responded to gentle prodding with a wire pick. The second group of worms, consisting of ∼2000 animals, was divided among three 6 cm pre-seeded RNAi + FUdR plates by chunking and were maintained at 20°C. At the sixth day of adulthood, ∼100 of these animals were subjected to heat stress according to the same procedure as outlined above for L4 thermal stress assays.

#### UV irradiation assay

Synchronized groups of worms were cultured on RNAi plates beginning at the L1 larval stage. Worms were maintained on those plates until the L4 larval stage when they were split into two groups. The first group of approximately 100 worms was distributed among three 3.5 cm pre-seeded RNAi + FUdR plates for UV treatment. Plates were placed in an irradiator (UV Stratalinker® 1800, Stratagene) with lids removed and were irradiated with 1mJ/m/s^2^ UV. Following treatment worms were maintained at 20°C and scored for survival daily until all animals had died. Worms were scored as alive if they responded to gentle prodding with a wire pick. The remaining second group of L4 larval worms were transferred to RNAi + FUdR plates by chunking and were maintained at 20°C until they reached Day 6 of adulthood when they were subjected to the same UV treatment and scoring regimen.

#### Fluorescence microscopy

Images of Plys-::GFP expression in *C. elegans* were taken at Day 6 on a Nikon Eclipse E800 equipped with a Jenoptik digital camera. Worms were immobilized on agar pads using polybead® polystyrene 0.10µm microspheres (Polysciences, Inc.), and pictures were taken quickly thereafter to avoid undo stress. Transgenic animals expressing *Plys7::gfp* were assigned to one of three categories based on the relative GFP expression levels (low, medium, and high) according to the following criteria: low—overall weak expression of GFP in the intestine or GFP expression limited to anterior of intestine, just behind the pharynx; medium—bright GFP expression from the anterior intestine to the midbody but little to no expression in posterior intestine; high—robust GFP expression throughout the entire length of intestine. The percentage GFP expression for each sample was calculated using a minimum of 200 animals and then the percentages were averaged across at least 3 biological replicates.

#### Statistical analyses

To calculate median lifespans (LT_50_s) a three parameter sigmoidal curve was fit to plots of survival (fraction of worms alive versus time) according to the general equation y=a/(1+e^(-(x-x0)/b)^) using SigmaPlot version 14 (Systat Software, San Jose, CA). This equation was used to determine the point at which 50% of the animals in the assay had died. The average fold difference between the LT_50_ of mutant strains or experimental RNAi treatments and control animals was calculated, and the statistical significance of that difference was assessed across all three replicates using a two-tailed Student’s t-test.

## Results

### *In silico* analysis of the C. elegans PP2A/4/6 phosphoprotein phosphatases

In both yeast and mammals significant structural and functional similarities between members of the PP2A/4/6 family of phosphoprotein phosphatases exist. In addition, proteomic and smaller scale biochemical studies have revealed non-canonical associations between subunits of the holoenzyme complexes (32,51,52). To ask how the PP2A/4/6 proteins in *C. elegans* compare to their human counterparts we first took an *in silico* approach. Alignments of the sequences of the three human catalytic subunits PP2Ac, PP4c and PP6c with their *C. elegans* orthologs revealed strong homology between them (Fig. S1, Table S1). For example, LET-92 and PPH-4.1 are both greater than 80% identical to PP2Ac and PP4c, respectively. Key structural motifs and regulatory residues are especially well-conserved both within catalytic subunits of the PP2A/4/6 family and across species. Based on these similarities, we anticipated that our studies would demonstrate significant parallels between these phosphatases in worms and humans.

We next asked whether *C. elegans* PP2A/4/6 proteins were expected to associate with the same binding partners as their human orthologs *in vivo*. To answer this question we performed database searches, using SMK-1 and PPH-4.1 as representative subunits for our analyses. We first used Wormbase to compile a list of PPH-4.1 interactors (Table S2). Drawing mainly from biochemically verified associations, high throughput yeast two-hybrid data, and interaction networks generated *in silico*, this list captures a broad spectrum of potential interactors. Included among them are not only proteins that could be members of a trimeric complex along with PPH-4.1 but also regulators, functional analogs, and possible substrates that may be dephosphorylated by PPH-4.1. In addition to SMK-1, two other worm homologs of PP4 regulatory subunits, PPFR-1 and PPFR-2, were included as potential PPH-4.1 interactors. The only other worm orthologue of a PP4 regulatory subunit, F46C5.6, was absent from the list of PPH-4.1 interactors found on Wormbase. Both orthologs of the yeast Tip41p-Tap42p regulatory system that modulates all members of the PP2A/4/6 family were identified as PPH-4.1 interactors, namely PPFR-4 and ZK688.9, orthologous to mammalian α4 and TIPRL, respectively. One other notable potential PPH-4.1 interactor is PPH-6, which encodes the *C. elegans* orthologue of PP6c, the catalytic subunit of the related but distinct PP6 complex. When we compared the list of potential PPH-4.1 interactors with the list of proteins predicted by Wormbase to interact with SMK-1, we expected to find significant overlap, especially among orthologues of the PP4 complex. Instead, although PPH-4.1 was listed as an interactor of SMK-1 the only other potential PP4 complex member that was in common to the list of interactors for both PPH-4.1 and SMK-1 was the regulatory subunit PPFR-1 (Table S3).

To gain further insight into the possible relationships between SMK-1, PPH-4.1, and their potential binding partners, we performed interaction network analysis using Genemania. All homologues of the PP2A/4/6 family earmarked as possible interactors of PPH-4.1 or SMK-1 on Wormbase along with the modulator PPFR-4 were included. This analysis yielded additional putative PPH-4.1 interactors (Fig S2). Among these were PAA-1 and Y71H2AM.20, *C. elegans* orthologues of human PR65α/PR65β and PTPA, respectively. This was surprising because in human cells these proteins associate with PP2c, the catalytic subunit of the PP2 complex, and not the PP4 complex. Specifically, PR65α/PR65β are scaffolding units of the PP2 complex, and PTPA has been shown to modify the catalytic site of the PP2c enzyme (53–55). This raises the intriguing possibility that certain subunits may associate with both a PP4c-containing complex and with PP2A complexes in *C. elegans*, a scenario that is supported by biochemical evidence in mammalian systems (32, 51).

Taken together, our database queries led to the unexpected finding that *C. elegans* orthologs of proteins not only of the PP4 complex but also of canonical subunits of the PP2A and PP6 holoenzyme complexes could potentially associate with SMK-1 and PPH-4.1. This lent credence to our experimental strategy of considering all *C. elegans* orthologs of the entire PP2A/4/6 family so that evidence supporting putative novel interactions between subunits in the context of aging would not be missed. Our database searches also revealed that evolutionarily conserved proteins that act to regulate the PP2A/4/6 holoenzymes by affecting the assembly of the complexes or by manipulating the active site of the catalytic subunit may also be binding partners of PPH-4.1.

Separate from posttranslational modifications of the PP2A/4/6 family catalytic subunits that influence their affinity for specific regulatory subunits, the catalytic activity of the enzymes is in parallel regulated by the evolutionarily conserved Tip41-Tap42 system. Constituents of this regulatory module may form transient or stable associations with PP2A/4/6 proteins. Originally identified in *Saccharomyces cerevisiae*, both the Tip 41 and Tap 42 proteins function in the TORC pathway (56, 57). Specifically, Tip41p antagonizes TORC by inhibiting Tap42, a negative regulator of the PP2A-related phosphatase Sit4p that dephosphorylates TORC substrates. In this manner Tip41p functions to activate PP2A-like catalysis by opposing Tap42. The mammalian ortholog of Tip41p, called TIPRL, on the other hand, cooperates with α4 (mammalian Tap42p) to promote mTORC activity by inhibiting the PP2A/4/6 phosphatases (58). In particular TIPRL and α4 together bind to holoenzymes and displace metal ions from the active site of the catalytic subunit to produce a latent yet stable complex. Catalytic activity of such decommissioned enzymes can be restored by PTPA, which reloads the metal ions back into position (59). *C. elegans* orthologs of all three of these proteins came up in our database searches for PPH-4.1 interactors. Since one of the objectives of our study was to uncover functional evidence that might implicate physical associations between proteins we therefore elected to include them as part of our functional analyses. In total we characterized 20 genes including each *C. elegans* ortholog of human PP2A/4/6 subunits along with orthologs of the regulatory proteins TIPRL, α4 and PPTA (Table 1).

**Table 1.**
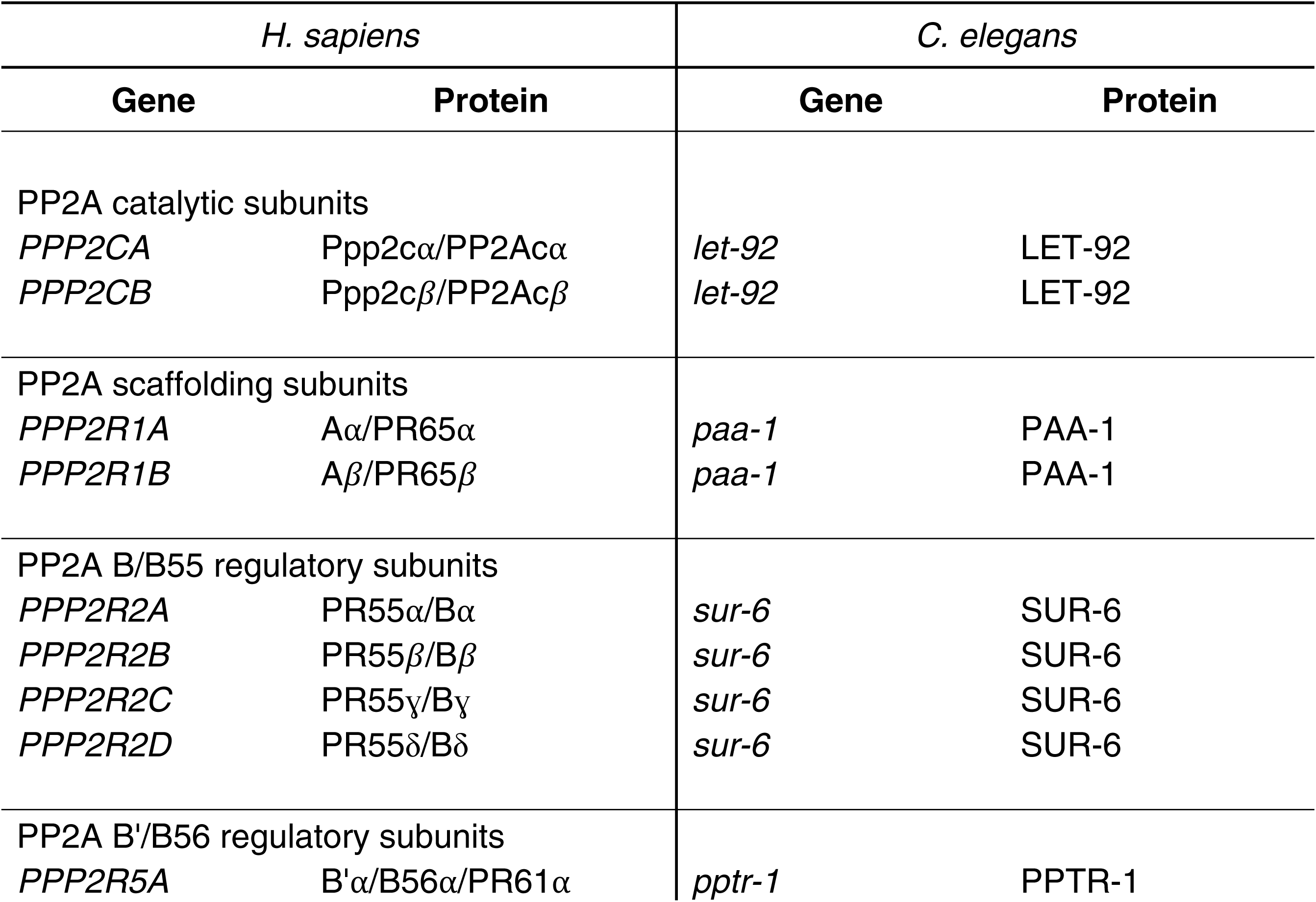
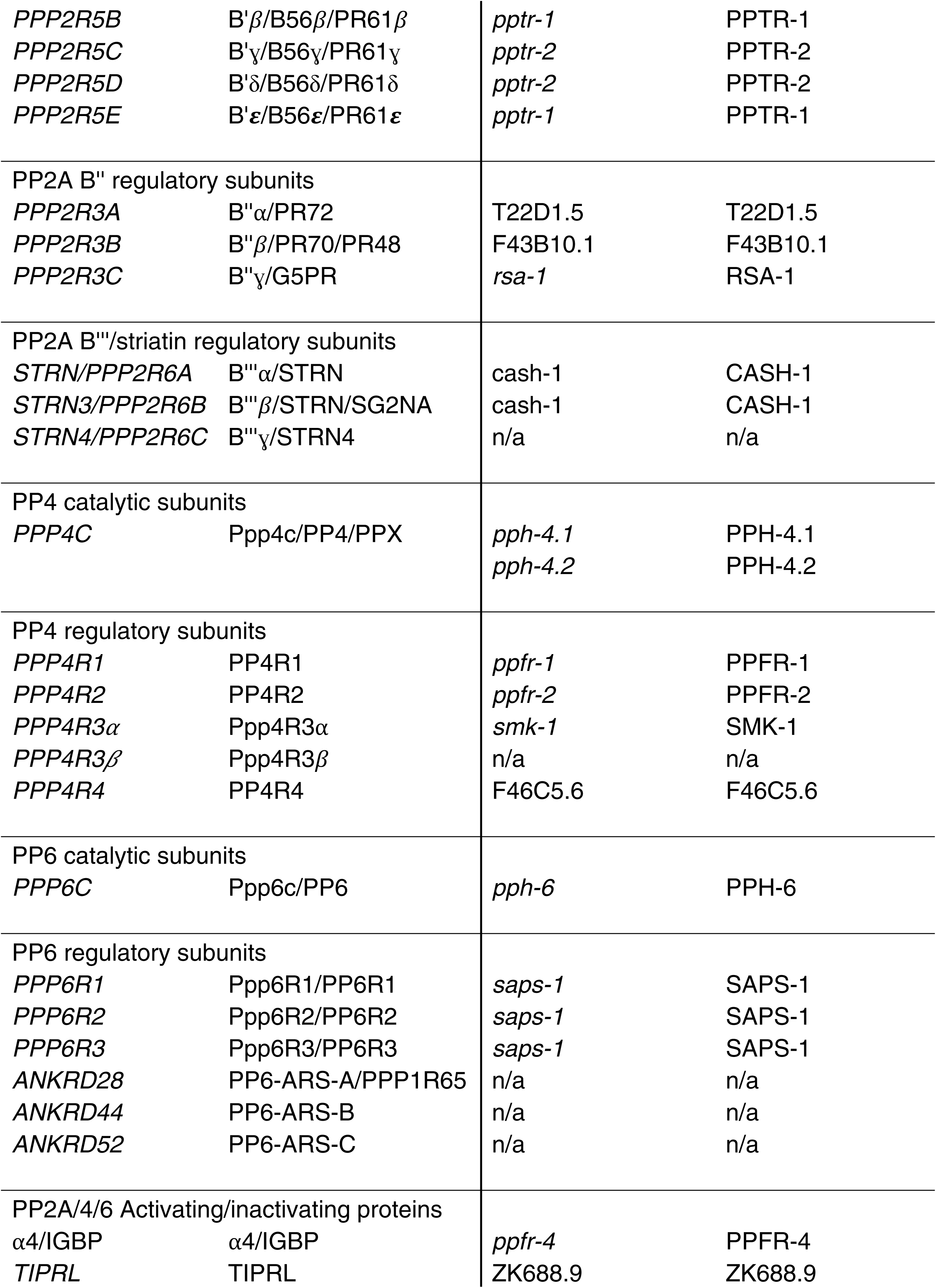
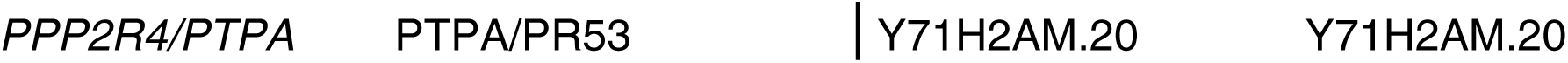
*C. elegans* orthologs of human PP2A/4/6 holoenzyme subunits and regulatory proteins.

The names of all human genes encoding catalytic, regulatory, or scaffolding constituents of PP2A/4/6 subfamily complexes are listed along with their orthologs in *C. elegans*. Genes encoding proteins that regulate the phosphatases through physical associations are also included. Corresponding protein names are provided.

#### Functional characterization of SMK-1 during aging

To investigate the PP2A/4/6 family of protein phosphatases during aging in *C. elegans* we began by studying the PP4 complex. We chose to focus on the SMK-1 regulatory subunit in particular as a test case to validate our experimental approach based on reverse genetics because direct binding between SMK-1 and the catalytic subunit PPH-4.1 had been demonstrated *in vitro* with recombinant proteins (44). In light of this connection, we expected that if SMK-1 functions as part of a PP4 complex in adult animals, RNAi knockdown of *smk-1*, *pph-4.1*, and potentially other constituent subunits of the complex should all result in similar if not identical phenotypes. Moreover, if a SMK-1/PPH-4.1-containing PP4 complex regulates DAF-16 during aging then RNAi targeting subunits of that complex should phenocopy RNAi targeting *daf-16*. Our strategy was to first compile a set of phenotypes attributable to *smk-1* inhibition in adult animals. These phenotypes would then generate a metric to which other genes encoding *C. elegans* orthologs of human PP4 complex members could be compared to determine the likelihood of them cooperating to produce the same output, including regulating the age-dependent increase in transcriptional activity of DAF-16. Further, functional evidence indicating that particular subunits act together would imply that they associate with each other *in vivo*. We anticipated that the same comparative analysis could be applied to putative constituents of the PP2A and PP6 complexes to determine whether they might also function during aging in *C. elegans*. Because we were interested in characterizing the SMK-1/PPH-4.1-containing complex in the context of aging, we conducted all of our studies on adult worms at Day 6 of adulthood when DAF-16 is transcriptionally active in parallel to control L4 larval stage animals (10). A two-tiered panel of assays was applied to evaluate the function of SMK-1 during aging. First, we asked whether SMK-1 is necessary for the transcriptional activity of DAF-16 in adult *C. elegans* by examining the expression of an *in vivo* reporter of DAF-16 activity, *Plys-7::GFP* in animals treated with RNAi targeting *smk-1*. We then functionally characterized *smk-1* by assessing its contribution to the ability of animals to resist acute environmental insults including bacterial infection with *Pseudomonas aeruginosa* (PA14), exposure to elevated temperature, and irradiation with ultraviolet (UV) light. In each stress assay, the phenotype resulting from *smk-1* knockdown was compared to the effect of knocking down *daf-16*.

We expected that if SMK-1 is necessary for the function of DAF-16 during adulthood, just as it is in the *daf-2(e1370)* mutant background, then RNAi targeting either *smk-1* or *daf-16* would have similar consequences. With minor exceptions, this is what we found. After initiating RNAi to knockdown *smk-1* or *daf-16* at the L1 larval stage, we first monitored the expression of the *Plys7::gfp* reporter over time by fluorescence microscopy (Fig. 1A). Compared to worms at the L4 larval stage, the expression of GFP driven by the promoter of *lys-7* in the intestines was substantially higher in Day 6 adults, consistent with an age-dependent increase in the transcriptional activity of DAF-16. In examining a population of animals we were able to assign individuals to one of three categories based on their GFP expression pattern (Fig. 1A). In general, we observed robust and uniform GFP expression along the length of the intestine among the majority of individuals in a synchronized population of Day 6 adults. These animals were scored as “high” GFP expressers. However, in a proportion of animals high levels of GFP expression were primarily confined to a few anterior intestinal cells with progressively weaker GFP expression in more posterior cells. Animals with this pattern of GFP expression were classified as “medium” GFP expressers. An even smaller fraction of worms expressed low levels of intestinal GFP that was frequently difficult to detect. RNAi targeting *daf-16* or *smk-1* had no effect on expression levels of the *Plys7::gfp* reporter in L4 larvae. By Day 6 of adulthood, however, *daf-16* knockdown resulted in a significant decrease in GFP transcription (Fig. 1B). This suggests that DAF-16 becomes activated in an age-dependent manner, consistent with previous studies (8, 9). We found that knocking down *smk-1* also resulted in decreased reporter expression in Day 6 adults (Fig. 1B), and so together our results indicate that the transcriptional activity of DAF-16 during aging in non-stressed wildtype animals may be regulated by SMK-1.

**Figure 1.**
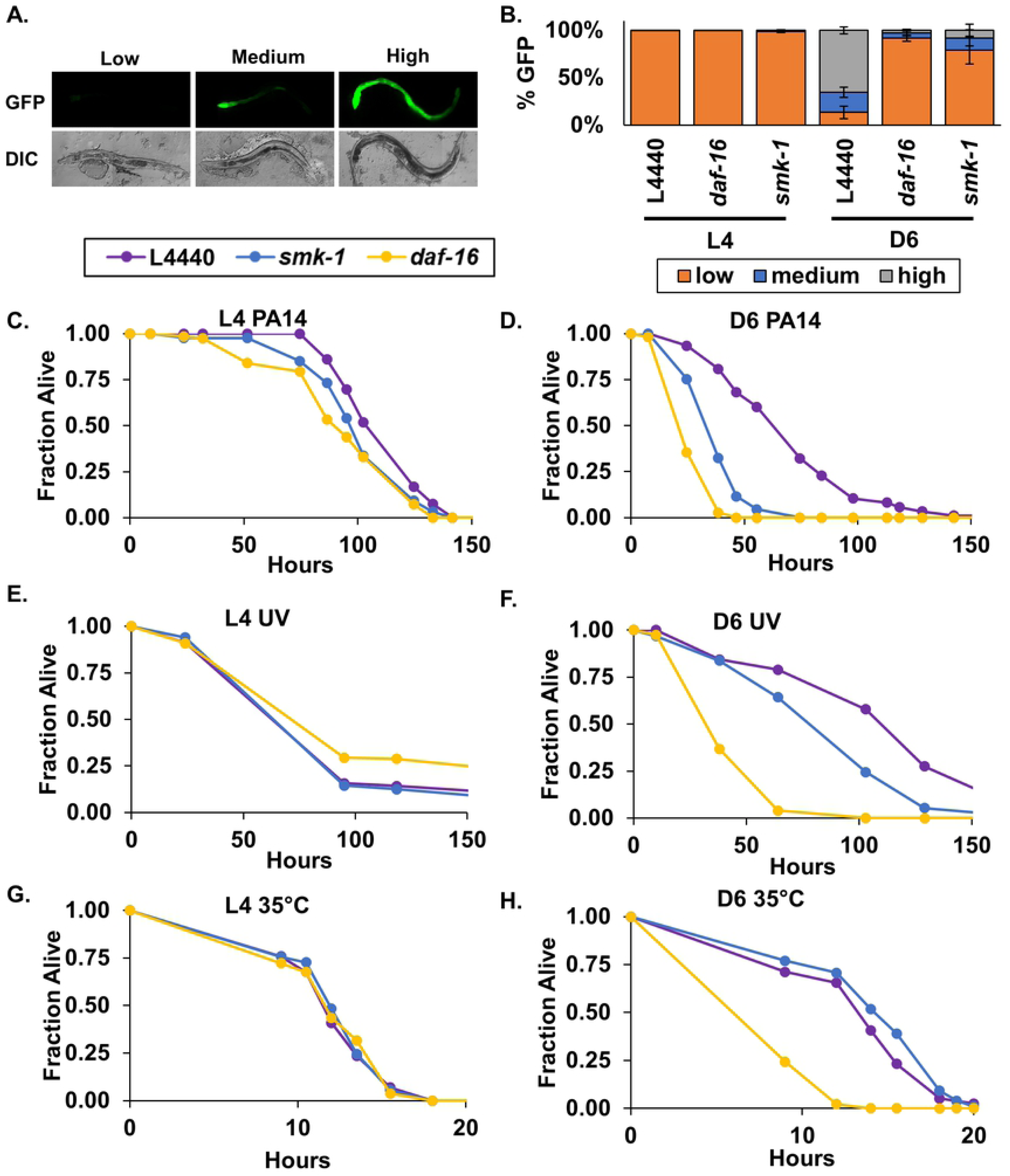
SMK-1 confers resistance to bacterial infection and ultraviolet radiation in adult *C. elegans.* (A) Animals expressing the *Plys7::gfp in vivo* reporter for DAF-16 transcriptional activity were examined by fluorescence microscopy at D6 of adulthood. Age-synchronized isogenic animals could be assigned to one of three categories based on the relative expression of GFP observed in the intestine. “Low”: little to no GFP expression “Medium”: GFP expression most apparent in the anterior portion of the intestine but also at lower levels toward the midbody. “High”: robust GFP expression along the entire length of the intestine. DIC, differential interference contrast image corresponding to the fluorescence image (GFP) shown above. (B) Quantification of worms in the Low (orange), Medium (blue), and High (grey) categories of *Plys7::gfp* expression at the L4 stage and D6 of adulthood following RNAi treatment targeting either *daf-16* or *smk-1*. n>200 animals for each RNAi treatment in three biological replicates. The average number of worms in each category is shown with error bars representing standard deviation from the mean. (C and D) Results of *Pseudomonas aeruginosa* infection assays. The survival of *C. elegans* subjected to RNAi against indicated genes beginning at L1 and exposed to *P. aeruginosa* continuously beginning at L4 (C) or D6 of adulthood (D) is plotted as the fraction of animals alive as a function of time. (E and F) Results of UV irradiation assays. Survival curves for *C. elegans* subjected to RNAi against the indicated genes from L1 until death and exposed to 1000mJ/s^2^ UV radiation at L4 (E) or D6 of adulthood (F). Hours represent time elapsed since the end of the irradiation treatment. (G and H) Results of heat stress assays. Survival curves for RNAi-treated *C. elegans* incubated at 35°C beginning at L4 (E) or D6 of adulthood (F) until they died. Hours represent the total time worms were incubated at 35° C. Beginning at the L1 stage, worms were maintained on RNAi plates without interruption for the duration of their lives. n>90 animals for each RNAi treatment in three biological replicates. Purple, L4440; yellow, *daf-16* RNAi; blue, yellow, *smk-1* RNAi.

Continuing to examine the functional parallels between the function of SMK-1 under low IIS conditions and during normal aging, we next asked whether SMK-1 is required to confer resistance to environmental stress in adult *C. elegans*. Consistent with the *Plys7::gfp* results, knocking down either *smk-1* or *daf-16* had no effect on the ability of L4 animals to resist bacterial infection, UV irradiation, or thermal stress (Fig. 1C, E, G; Table 2). On the other hand, both *smk-1* and *daf-16* were necessary for Day 6 adults to withstand challenge with either *P. aeruginosa* infection (Fig. 1D) or with UV irradiation (Fig. 1F), as knockdown of either gene reduced the median survival of animals exposed to these insults (Table 2). We found that *daf-16* but not *smk-1* contributes to resistance from thermal stress in adult *C. elegans*, consistent with the role of SMK-1 in *daf-2(e1370)* mutants (Fig. 1 G, H) (43, 45). Taken together the results of our stress assays demonstrate a cooperation between SMK-1 and DAF-16 in wild type animals. They suggest that during aging SMK-1 modulates the transcriptional activity of DAF-16, thereby regulating DAF-16-mediated resistance to multiple insults including bacterial infection and ultraviolet irradiation.

**Table 2.**
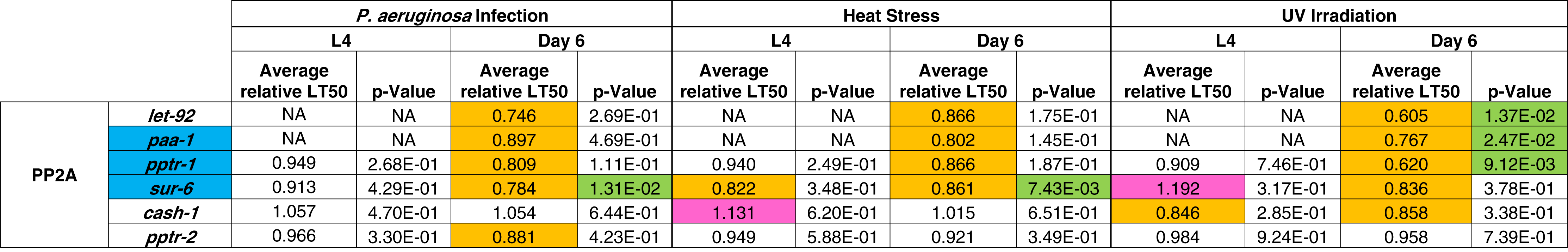
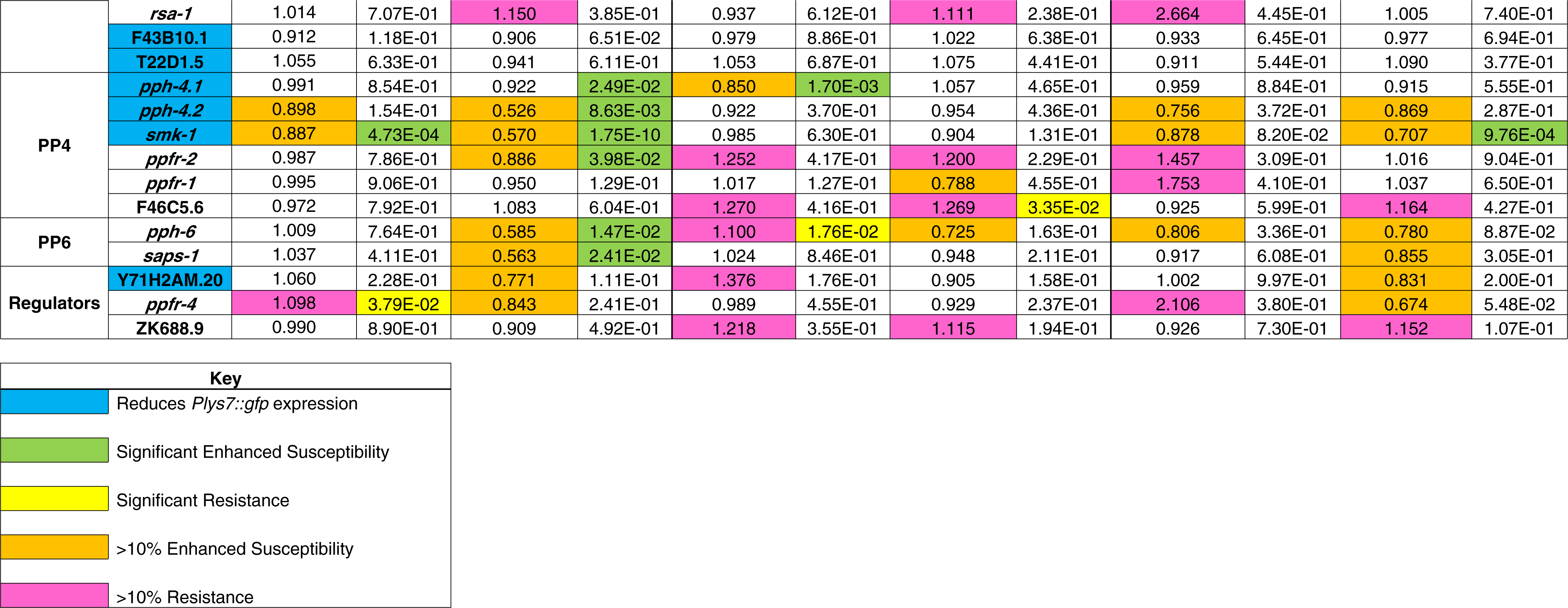
Summary of phenotypes resulting from knockdown of members of the PP2A/4/6 family and their regulators.

Genes tested in our functional analyses are grouped according to the PP2A/4/6 holoenzyme with which they are typically associated. Orthologs of regulators of the PP2A/4/6 family in humans are grouped separately. The table presents aggregated data of phenotypes observed for RNAi-treated animals in the DAF-16 reporter expression assay and in each of the three stress assays. Genes whose names are highlighted in blue are required for the increase in *Plys-7::GFP* expression at Day 6. Each row lists the average relative median lifespan (relative LT_50_) for L4 and Day 6 adults challenged with the indicated environmental stress upon knockdown of one particular gene. Average relative LT_50_ was calculated by dividing the average LT_50_ of RNAi-treated animals in a given stress assay by the average LT_50_ of age-matched control animals in that same assay. Therefore, it represents the proportional increase or decrease in median lifespan under stressful conditions that are attributable to RNAi inhibition of the gene of interest. Where knockdowns enhanced sensitivity to stress, decreasing median lifespan after exposure to stress by more than 10%, the average relative LT_50_ value is shaded in orange. When RNAi treatment conferred resistance to stress and increased the median lifespan of stressed worms by more than 10%, the average relative LT_50_ value is shaded in pink. Statistical significance was determined by Student’s t-test, with a threshold of p<0.05. p-values associated with significant susceptibility and resistance are shaded in green and yellow, respectively.

#### Putative subunits of the PP4 and PP2A complexes influence the transcriptional activity of DAF-16 in adult *C. elegans*

To expand our analyses to include all putative subunits of the PP4 complex and, in fact, to the entire PP2A/4/6 family, we first surveyed the effect of knocking down each gene individually on the age-dependent increase in *Plys7::gfp* expression (Fig. 2). RNAi treatments that resulted in greater than 25% of worms being assigned to the low GFP expression category were considered to function in a similar manner to SMK-1 to regulate the activity of DAF-16. None of the genes that we tested had a significant effect on the expression of the *Plys7::gfp* reporter in L4 larvae (Fig. 2A). At Day 6, knockdown of the PP4 catalytic subunit *pph-4.1* and its paralog *pph-4.2* had some of the strongest effects on GFP expression, resulting in a higher proportion of worms in the low expressing category, similar to the effect of knocking down *smk-1* (Fig 2B). This result was expected since SMK-1 and PPH-4.1/PPH-4.2 associate *in vivo* (44, 45). RNAi targeting F46C5.6 and *ppfr-2*, *C. elegans* homologues of two other PP4 regulatory subunits, also modestly increased the proportion of worms expressing low levels of GFP, yet this effect was not statistically significant. Notably, inhibiting the expression of several worm homologues of PP2 complex members also suppressed the age-dependent increase in *plys7::gfp* expression. Inhibition of *pptr-1*, *sur-6*, T22D1.5, *paa-1*, and F43B10.1 all resulted in an increase in low levels of GFP expression at Day 6 relative to control animals. In addition, Y71H2AM.20, the *C. elegans* ortholog of the regulatory protein PTPA was also found to be necessary for the age-dependent increase in *plys-7::GFP*. These results suggest that SMK-1 and PPH-4.1 function together in adult *C. elegans*, and they implicate roles for both the PP4 and PP2A complexes in regulating the transcriptional activity of DAF-16 during aging.

**Figure 2.**
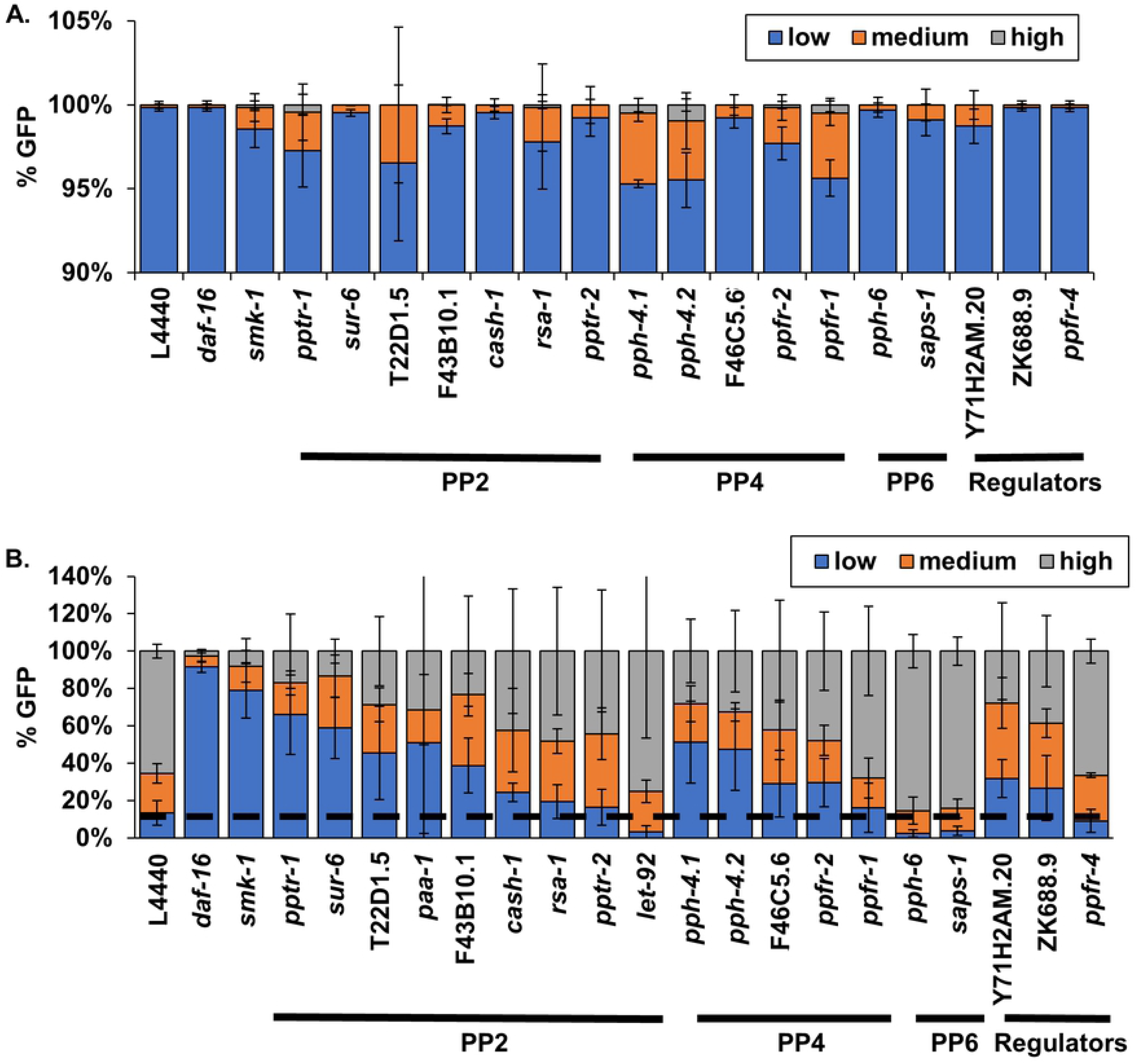
Genes encoding putative members of the PP2A, PP4, and PP6 complexes are required for the age-dependent increase in DAF-16 transcriptional activity. Quantification of worms in the Low (orange), Medium (blue), and High (grey) categories of *Plys7::gfp* expression at the L4 stage (A) and D6 of adulthood (B) following RNAi treatment targeting the indicated gene. n>200 animals for each RNAi treatment in three biological replicates. The average number of worms in each category is shown with error bars representing standard deviation from the mean.

#### Members of the PP2A/4/6 subfamily contribute to innate immunity during aging

We found that *smk-1* is necessary to confer resistance to bacterial infection in Day 6 adults, as is DAF-16 (Fig 1D). This result leads to the prediction that *pph-4.1*, encoding the catalytic subunit of the PP4 complex, is also required for host defense during aging. Further, based on the results of our *in vivo* reporter assay for DAF-16 transcriptional activity, we wondered whether PP2A subunits may also function in innate immunity later in life. We addressed these possibilities as part of our characterization of the PP2A/4/6 family in adults challenged with *P. aeruginosa*. In a phenotype reminiscent of the *smk-1* knockdown, RNAi targeting homologues of the PP4 catalytic subunit *pph-4.1* or its paralog *pph-4.2* reduced the ability of Day 6 adults to resist bacterial infection but had no effect on the susceptibility of L4 larvae to pathogen (Fig. 3A, B, G, H). The magnitude of the resulting decrease in median lifespan (LT_50_) of infected animals caused by inhibiting *pph-4.2* expression was comparable to the effect of *smk-1* knockdown, but *RNAi* targeting *pph-4.1* had a considerably milder effect across our replicates (Table 2). Coupled with the results of our *in vivo* reporter assay, this result represents a second independent circumstance in which RNAi inhibition of *smk-1* phenocopied the knockdown of *pph-4.1/4.2*. We also considered it as providing validation to our experimental approach since our functional analyses confirmed biochemically verified interactions between SMK-1 and PPH-4.1/4.2.

**Figure 3.**
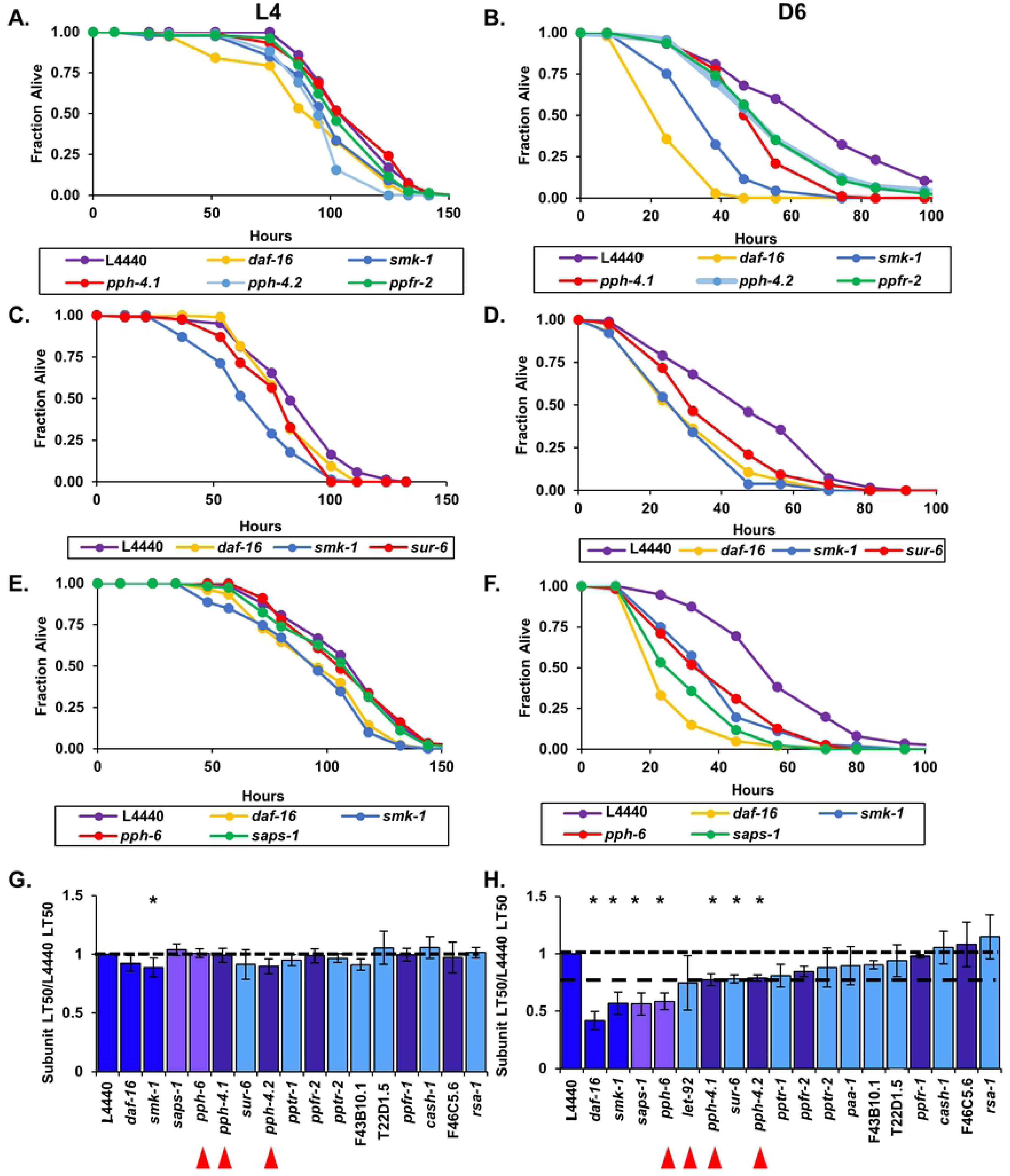
Members of putative PP4 and PP6 holoenzymes along with the PP2A regulatory subunit SUR-6 are required for innate immunity during adulthood in *C. elegans*. Following RNAi treatment beginning at the L1 stage to target *C. elegans* homologs of catalytic and regulatory subunits of the PP4 (A and B), PP2A (C and D), and PP6 (E and F) complexes, worms were infected with *P. aeruginosa* at the L4 larval stage (A,C,E) or at D6 of adulthood (B,D,F). The fraction of worms alive at each time point after infection was initiated is plotted as a function of time in hours. In all cases RNAi targeting *daf-16* or *smk-1* and the empty RNAi vector L4440 were included as controls. Only RNAi knockdowns that produced statistically significant phenotypes are shown. (G and H) The average median survival (LT_50_) of animals treated with RNAi targeting the indicated genes following infection with *P. aeruginosa* at L4 (G) or D6 (H) is shown as a fraction of the average median survival of L4440 controls. Bars, standard error of the mean (SEM). Bar colors correspond to the protein phosphatase complex to which products of the indicated genes belong or to controls. Dark blue: L4440, *daf-16*, and *smk-1*; light blue: PP2A; dark purple: PP4; light purple: PP6. Asterisks indicate RNAi treatments producing statistically significant differences in median survival (p<0.05). Horizontal lines are drawn at a relative median survival of 1 and, for a reference in (H) the relative median survival of adult worms treated with RNAi against *pph-4.1*. Red arrowheads are beneath the names of genes encoding catalytic subunits of the PP2A, 4 and 6 complexes.

Along with *pph-4.1/4.2* and *smk-1*, RNAi inhibition of another PP4 regulatory subunit homologue, *ppfr-2* (PP4R2 in humans), also reduced the median lifespan of worms infected at Day 6 of adulthood but had no effect on the ability of L4 larvae to resist infection (Fig. 3A, B, G, H). No other PP4 regulatory subunits were found to be required for innate immunity in Day 6 adults (Fig. 3H; Fig S3). These results imply that PPFR-2 may act in conjunction with SMK-1 and PPH-4.1 to protect adult *C. elegans* from bacterial pathogens.

We found genes encoding putative subunits of other protein phosphatase complexes to also be important for host defense in adult *C. elegans*. In addition to the effect of inhibiting the expression of PP4 subunits, one of the ten RNAi treatments directed against *C. elegans* orthologs of PP2A subunits enhanced the susceptibility of adult animals to *P. aeruginosa* infection. Knockdown of *sur-6*, the sole worm homolog of human PP2A B/B55 regulatory subunits, significantly reduced the median lifespan of infected Day 6 adults but had no effect on larvae (Fig. 3C, D, G, H; Table 2). We found no evidence of a role for other orthologs of PP2A complex subunits in adult or larval innate immunity (Fig. 3H; Fig. S4). In an unexpected result, inhibiting the expression of two orthologues of PP6 complex members *pph-6*, the catalytic subunit, and *saps-1*, a regulatory subunit, accelerated the rate of death from infection when worms were challenged with *P. aeruginosa* at Day 6 of adulthood but not at the L4 larval stage (Fig. 3E, F, G, H). This was surprising because none of the PP6 members appear to be required for the increased expression of *plys7::gfp* in adults, a departure from *smk-1*-like phenotypes (Fig. 2B). Taken together, our results indicate that the entire PP2A/4/6 family plays a role in innate immunity during aging and that the PP6 complex may do so without influencing the transcriptional output of DAF-16.

#### SMK-1 and the PP2A complex confer UV resistance in postreproductive *C. elegans*

We found SMK-1 to be important not only for host defense in Day 6 animals, but also for resistance to ultraviolet irradiation, consistent with its role in *daf-2(e1370)* mutants (Fig 1F; Table 2) (43, 60). We therefore expected that other components of the PP4 complex would be required for survival upon exposure to UV light during aging. This was not the case. No canonical members of the PP4 holoenzyme, including the catalytic subunits, seem to contribute to UV tolerance in *C. elegans* as knocking them down did not affect the survival of irradiated L4 larvae or Day 6 adults (Fig. 4A, B, G, H; Fig. S5). On the other hand, inhibiting expression of the PP2A catalytic subunit, *let-92*, the scaffolding protein *paa-1* and the regulatory subunit *pptr-1* caused Day 6 adult animals exposed to UV light to die more rapidly, leading to significant reductions in their median lifespans (Fig. 4 C, D, G, H; Table 2). Other orthologs of PP2A subunits do not appear to play a role in conferring resistance to UV light (Fig. S6). Targeting PP6 complex subunits by RNAi was also inconsequential to the ability of adults to resist UV irradiation (Fig. 4E, F, G, H; Fig. S7). Our data therefore suggest that a complete PP2A holoenzyme is involved in the ability of post-reproductive animals to resist UV exposure and that it may act in conjunction with SMK-1.

**Figure 4.**
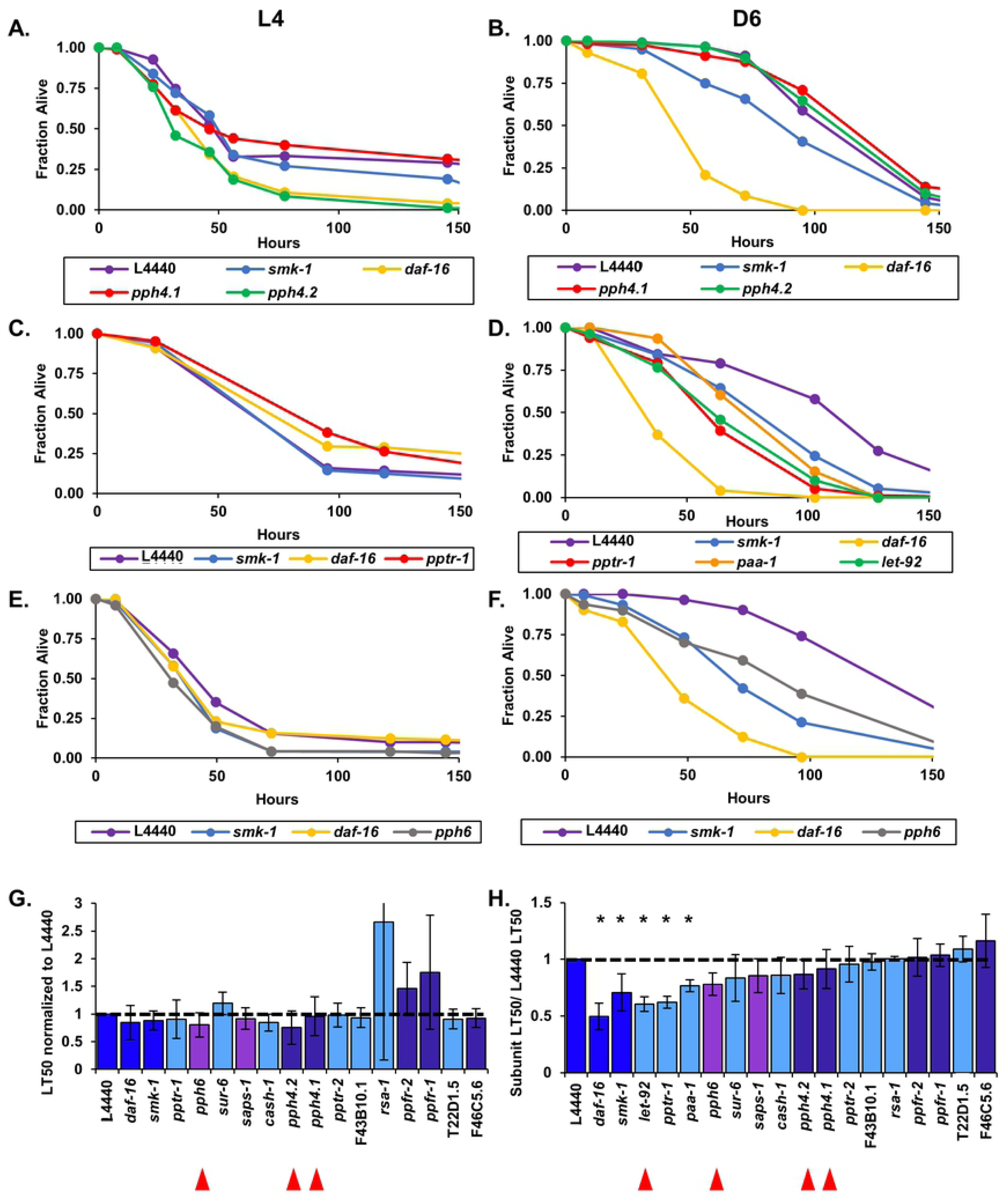
Resistance to UV irradiation in adult *C. elegans* is conferred by the PP2A complex and SMK-1. RNAi treatment was initiated at the L1 stage to target *C. elegans* homologs of catalytic and regulatory subunits of the PP4 (A and B), PP2A (C and D), and PP6 (E and F) complexes and was continued for the duration of the assay. Worms were exposed to UV radiation at the L4 larval stage (A,C,E) or at D6 of adulthood (B,D,F) after which their survival under standard culturing conditions was monitored. The fraction of worms alive at each time point following the UV treatment is plotted as a function of time. In all cases RNAi targeting *daf-16* or *smk-1* and the empty RNAi vector L4440 were included as controls. Only RNAi knockdowns that produced statistically significant phenotypes are shown. (G and H) The average median survival (LT_50_) of animals treated with RNAi targeting the indicated genes following exposure to UV light at L4 (G) or D6 (H) is shown as a fraction of the average median survival of L4440 controls. Bars, standard error of the mean (SEM). Bar colors correspond to the protein phosphatase complex to which products of the indicated genes belong or to controls. Dark blue: L4440, *daf-16*, and *smk-1*; light blue: PP2A; dark purple: PP4; light purple: PP6. Asterisks indicate RNAi treatments producing statistically significant differences in median survival (p<0.05). Horizontal lines are drawn at a relative median survival of 1. Red arrowheads are beneath the names of genes encoding catalytic subunits of the PP2A, 4 and 6 complexes.

#### The PP2A regulatory subunit SUR-6 promotes resistance to heat stress in adult worms

Despite its roles in conferring resistance to bacterial pathogens and ultraviolet irradiation, SMK-1 does not contribute to thermotolerance in adult *C. elegans* even though DAF-16 does (Fig. 1G, H). In like manner, we found no evidence to suggest that the PP4 catalytic subunits PPH-4.1 or PPH-4.2 function to protect animals from thermal stress during aging (Fig. 5C, D, G. H). However, RNAi targeting a putative regulatory subunit F46C5.6, the ortholog of PP4R4, resulted in a surprising modest but significant increase in resistance to high temperatures in Day 6 adult worms but not larvae (Fig. 5A, B, G, H). Neither of the other two *C. elegans* orthologs of PP4 regulatory subunits PPFR-1 and PPFR-2 appear to influence survival of heat stressed animals at the larval or adult stages (Fig. 5G, H; Fig. S8). On its own this result suggests that in contrast to the roles of other PP4 regulatory subunits, F46C5.6 could function to inhibit the catalytic activity of the PP4 complex in adult animals. Yet considering that there is no corresponding enhanced sensitivity to heat stress upon knockdown of *pph-4.1* or *pph-4.2*, this scenario seems unlikely.

**Figure 5.**
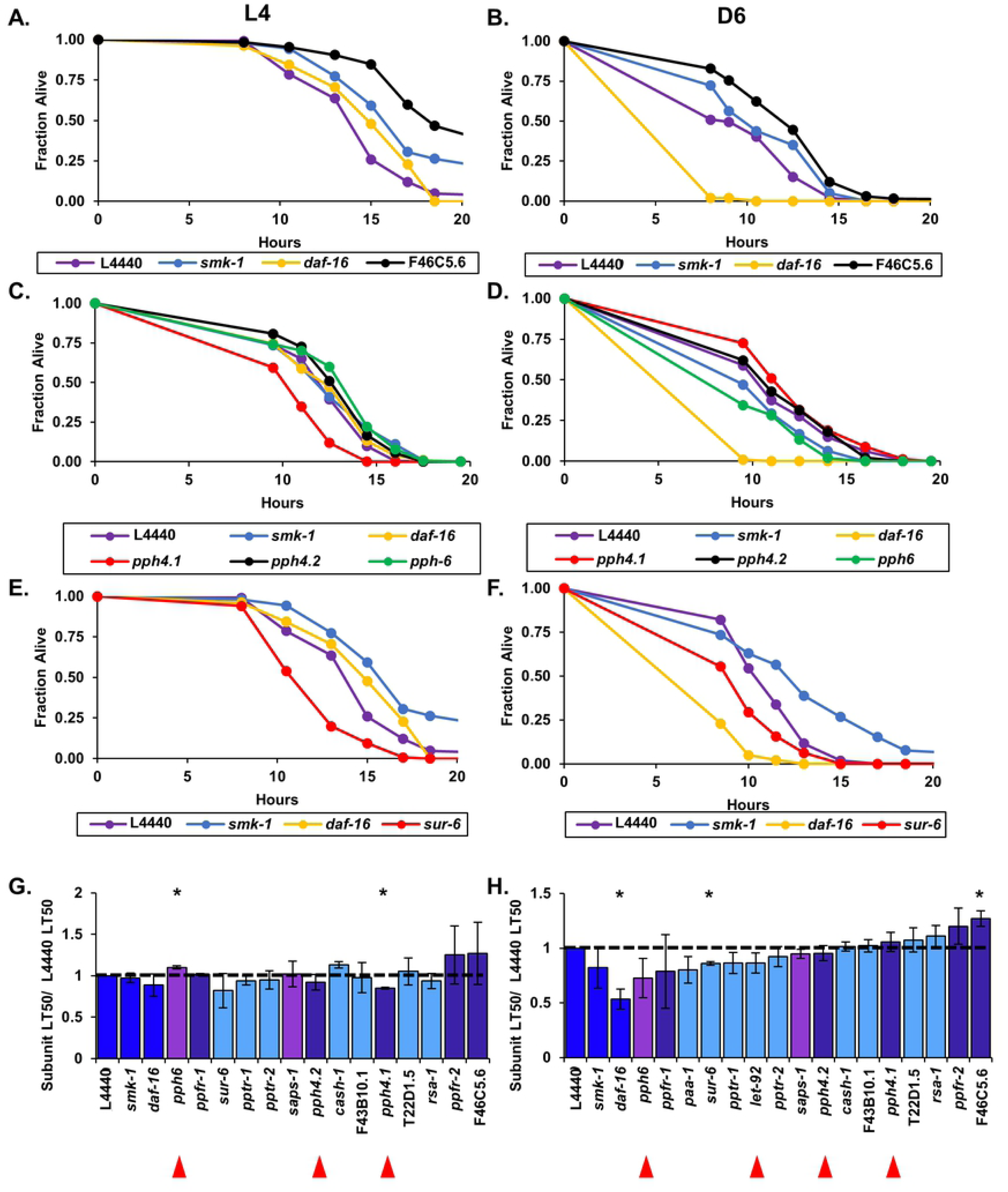
The PP2A subunit SUR-6 contributes to thermotolerance in adult worms. RNAi treatment targeting *C. elegans* homologs of catalytic and regulatory subunits of the PP4 (A-D), PP6 (C and D), and PP2A (E and F) complexes was initiated at the L1 stage and continued for the duration of the assay. To induce thermal stress at the L4 larval stage (A,C,E) or at D6 of adulthood (B,D,F) worms were shifted from 20° C to 35° C until all of the animals had died. The fraction of worms alive at each time point during the incubation at high temperature is plotted as a function of time in hours. In all cases RNAi targeting *daf-16* or *smk-1* and the empty RNAi vector L4440 were included as controls. Only RNAi knockdowns that produced statistically significant phenotypes are shown. (G and H) The average median survival (LT_50_) of animals treated with RNAi targeting the indicated genes following the shift to 35° C at L4 (G) or D6 (H) is shown as a fraction of the average median survival of L4440 controls. Bars, standard error of the mean (SEM). Bar colors correspond to the protein phosphatase complex to which products of the indicated genes belong or to controls. Dark blue: L4440, *daf-16*, and *smk-1*; light blue: PP2A; dark purple: PP4; light purple: PP6. Asterisks indicate RNAi treatments producing statistically significant differences in median survival (p<0.05). Horizontal lines are drawn at a relative median survival of 1. Red arrowheads are beneath the names of genes encoding catalytic subunits of the PP2A, 4 and 6 complexes.

Out of all of the other members of the PP2A/4/6 family that we tested, only RNAi targeting *sur-6*, encoding a regulatory subunit of PP2A, reduced the survival of Day 6 adult *C. elegans* maintained at elevated temperature (Fig 5E, F, G, H). Knockdown of orthologs of the catalytic, regulatory, or scaffolding subunits of PP2A did not affect the survival of adults challenged by thermal stress (Fig. 5H; Fig. S9). Although inhibiting *pph-6* expression increased the resistance of L4 worms the thermal stress (Fig. 5C), this effect did not persist into adulthood (Fig. 5D). Upon exposure to high temperature, the survival of Day 6 animals treated with RNAi targeting *pph-6* or *saps-1* was comparable to the control group, (Fig. 5H; Fig. S10). Our do not implicate a complete PP2A/4/6 holoenzyme as functioning to confer resistance to heat stress during adulthood, However, individual members of the PP2A/4/6 family may influence the response to thermal stress as part of other pathways in an age-specific manner.

#### Characterization of putative regulators of the PP2A/4/6 family that are not incorporated into the complexes as canonical subunits

In mammals one group of regulatory proteins modulates the function of the entire PP2A/4/6 subfamily of protein phosphatases through mechanisms involving physical interactions and not post-translational modifications. An example of such a protein is α4, which protects the catalytic subunits of the PP2, PP4, and PP6 complexes in humans by inhibiting their ubiquitin-mediated degradation (61). At the same time, α4 can act as a negative regulator by functioning together with TIPRL. For example, α4 and TIPRL bind noncompetitively to PP2Ac and inactivate it by both expelling metal ions from the active site and displacing the scaffold and regulatory subunits (58). Conversely, PTPA can reactivate PP2A/4/6 phosphatases by reloading metal ions back in to the active site (59). Although they are not considered to be constituents of the PP4 holoenzyme, orthologs of all three of these regulators were identified in our database queries as potential interactors of SMK-1/PPH-4.1 (Fig. S2; Table S2). We therefore asked whether they might modulate the PP2A/4/6 family during aging in *C. elegans*. Since our data indicate that the PP2A/4/6 family is important for resistance to environmental stress in adult animals, we expected that if PPFR-4 and ZK688.9 cooperate to inhibit the phosphatases in *C. elegans* as they do in mammals, then knocking them down would increase the survival of Day 6 adults challenged with acute insults. Further, since PTPA acts antagonistically to α4 and TIPRL in mammals, we anticipated that RNAi targeting its ortholog Y71H2AM.20 would cause Day 6 adults to become more sensitive to stress. While knocking down Y71H2AM.20 did increase the susceptibility of Day 6 adults to bacterial infection and their sensitivity to UV light, neither effect was statistically significant (Table 2; Fig. 6E, F; Fig. S11; Fig. S12). We found no evidence of a role for Y71H2AM.20 in protecting animals from thermal stress (Fig. 6G; Fig. S13). Surprisingly, instead of making adult animals more resistant, when *ppfr-4* was targeted by RNAi, Day 6 *C. elegans* were more sensitive to UV irradiation, yet the effect on the relative median survival of these animals fell just short of the significance threshold (p= 0.0548) (Fig. 6A, B, E; Table 2). The only resistance phenotype that we found to be associated with *ppfr-4* knockdown was when L4 animals were challenged with *P. aeruginosa* but not when adults were infected (Fig. 6 C, D, F). Our observations suggest that there may be some differences between the roles of this class of PP2A/4/6 regulators in *C. elegans* and mammals, but the lack of statistical significance in our data prevents us from drawing definitive conclusions.

**Figure 6.**
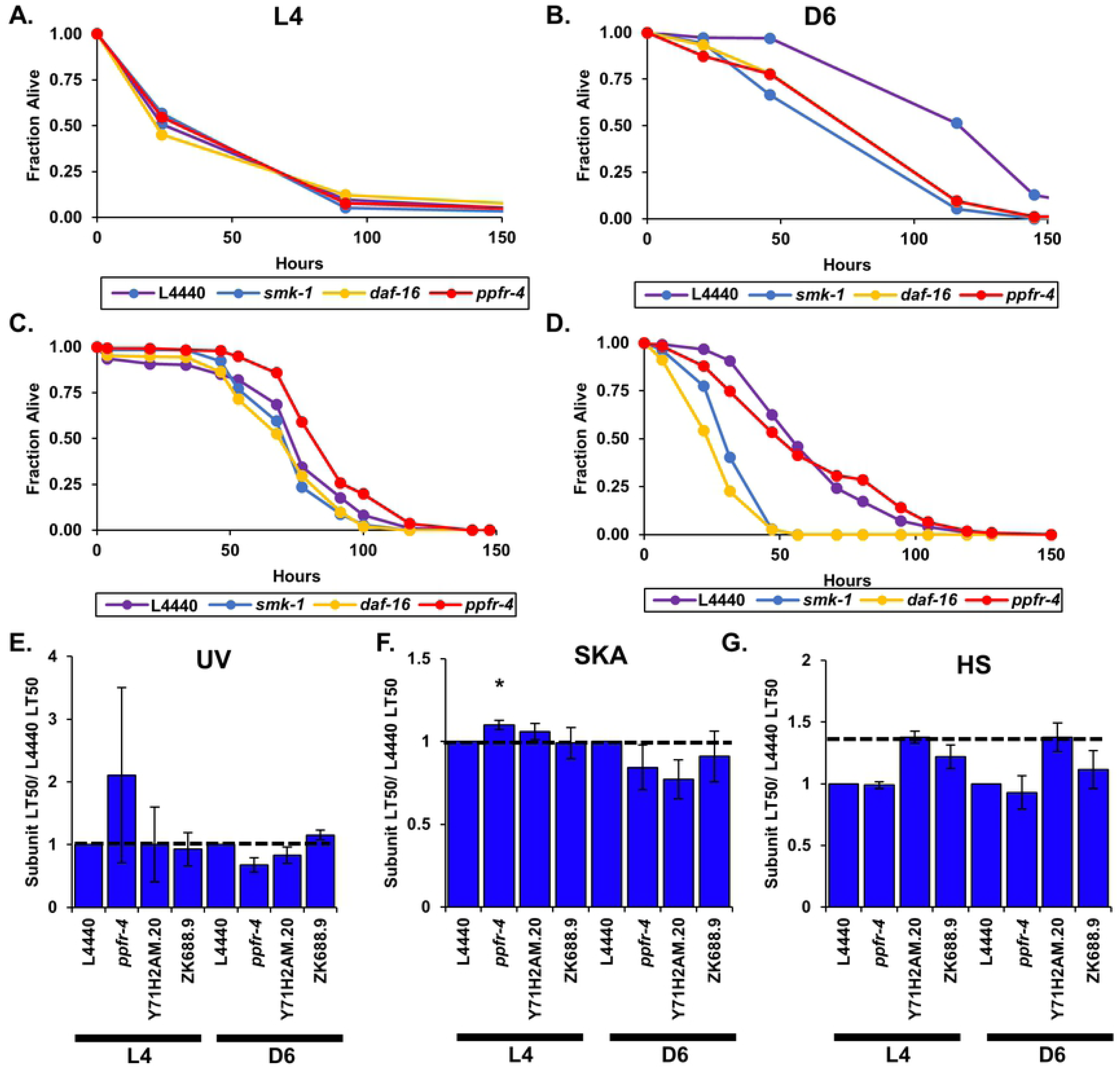
PPFR-4, the *C. elegans* ortholog of the α4 regulatory protein, may be required for resistance to UV radiation during adulthood. (A-D) Beginning at the L1 larval stage worms were treated with RNAi to knockdown *ppfr-4*. At the L4 stage (A) or Day 6 of adulthood (B) worms were irradiated with UV light and then returned to standard culture conditions where their survival was monitored over time. The fraction of worms alive at each time point following the UV treatment is plotted as a function of time. RNAi targeting *daf-16* or *smk-1* and the empty RNAi vector L4440 were included as controls. At the L4 stage (C) or Day 6 of adulthood (D) worms were transferred to plates seeded with *P. aeruginosa* where their survival was monitored over time. The fraction of worms alive at each time point following the UV treatment is plotted as a function of time. RNAi targeting *daf-16* or *smk-1* and the empty RNAi vector L4440 were included as controls. (E-G) The average median survival (LT_50_) of L4 and D6 animals treated with RNAi targeting the indicated genes following exposure to UV light (E), infection with *P. aeruginosa* (F) or incubation at 35° C (G) is shown as a fraction of the average median survival of L4440 controls. Bars, standard error of the mean (SEM). Asterisks indicate RNAi treatments producing statistically significant differences in median survival (p<0.05). Horizontal lines are drawn at a relative median survival of 1. UV, ultraviolet light; SKA, slow kill assay (*P. aeruginosa* infection); HS, heat stress (35° C).

## Discussion

The FoxO transcription factor DAF-16 is a major longevity determinant in *C. elegans*. Animals with constitutively active DAF-16 live longer and are more youthful than their wildtype counterparts, suggesting that DAF-16 contributes not only to lifespan but also to healthspan (62). Accordingly, one category of DAF-16 transcriptional targets are genes that confer resistance to stress, thus shielding animals from long-lasting cellular damage (7). While exposure to environmental insults induces DAF-16 to upregulate these genes, recent evidence suggests that DAF-16 becomes activated in age-dependent manner in animals that have not been challenged with a stressor (8, 9). Here we investigated the mechanism by which DAF-16 is activated during aging in *C. elegans*. We focused our studies on the PP2A/4/6 family of phosphoprotein phosphatases, which have been implicated in regulating FoxO transcription factors in evolutionarily diverse species (14–17,19,21,45). Using a reverse genetics approach, we performed functional analyses to survey the complete set of *C. elegans* orthologs of human PP2A/4/6 regulatory and catalytic subunits in the context of aging. Our results suggest that the transcriptional activity of DAF-16 in adult animals is regulated by both PP2A and PP4 complexes. The specific effect of these complexes on DAF-16 appears to be non-overlapping, as each is important for a particular facet of DAF-16-mediated stress resistance. In the course of our studies, we also uncovered evidence of an apparent DAF-16-independent role for the PP6 phosphatase in host defense in post-reproductive adult *C. elegans*, thus highlighting the importance of the entire PP2A/4/6 family during aging.

The PP2A/4/6 family of phosphoprotein phosphatases regulates a broad spectrum of processes essential to the survival and normal physiology of a variety of cell types. This is achieved in part by controlling pathways that are foundational to higher order cellular function. For example, both the biogenesis and the mRNA splicing function of small nuclear ribonucleoproteins (snRNPs) are regulated by PP4 and PP6, while PP2A contributes to mRNA surveillance through nonsense-mediated decay (NMD) (63–65). PP2A/4/6 phosphatases also regulate membrane trafficking and influence the dynamics and metabolism of mitochondria (66, 67). Further, the activity of key components of major signaling pathways, including NF-KB and JNK, is subject to regulation by PP2A/4/6 (35,68–76). This particular group of phosphatases is perhaps best known, however, for the key roles that its members play with regard to the cell cycle. One of the most critical functions of the PP2A/4/6 family is to negatively regulate the kinases that drive progression of the cell cycle (30,37,38,40,42,76,77). Across evolutionarily diverse species, when cells are permitted to divide the PP2A/4/6 phosphatases ensure the faithful transmission of genetic material to daughter cells by regulating the assembly and positioning of the spindle, the cohesion of chromatin, and the attachment of microtubules to kinetochores (30,78–86). Both in the context of mitosis and during interphase PP4 and PP6 in particular further contribute to genome stability by promoting DNA repair and by resolving changes in chromatin structure associated with the response to DNA damage (87–97). Given these functions, it is perhaps not surprising that members of the PP2A/4/6 family have been identified as tumor suppressors (98–102). At the whole organism level, the PP2A/4/6 family is especially important at the very early stages of life, ensuring the survival and developmental progression of embryos (36,103,104). In addition, these phosphatases orchestrate the development of multiple tissues including bone, adipose, innate immune cells, and neurons (34–36,105,106). Certain other functions of PP2A/4/6 indicate that later in life they contribute to longevity. For example, PP6 suppresses inflammation and can promote autophagy, both of which would preserve normal lifespan (107, 108). Regulation of IIS components by PP2A is an example of an even more direct connection between the PP2A/4/6 family and aging, since insulin signaling is a predominant longevity determinant in invertebrates and other animals (1). Considering these observations and in light of the breadth and magnitude of their impact on cellular function, we were particularly interested in investigating the function of the PP2A/4/6 phosphatases during aging, a context in which they have not yet been extensively studied. We asked whether the PP2A/4/6 phosphatases might function in adult *C. elegans* to regulate the age-dependent increase in DAF-16 transcriptional activity and contribute to stress resistance.

To investigate the potential role of the PP2A/4/6 phosphatases in regulating DAF-16 during aging in *C. elegans*, we used a two-pronged approach where in each branch the phenotype resulting from RNAi knockdown of candidate genes was always compared to the phenotype associated with reduced DAF-16 expression. First we asked whether RNAi-mediated knockdown of genes encoding worm orthologs of human PP2A/4/6 subunits reduced the age-dependent increase in the expression of *plys-7::GFP*, an in vivo reporter of DAF-16 transcriptional activity. We then challenged RNAi-treated adult animals with three different types of stress to which DAF-16 is known to confer resistance—bacterial infection, heat, and ultraviolet irradiation. Based on this analysis, we identified 9 members of the PP2A/4/6 family that recapitulated *daf-16* knockdown phenotypes in at least one of the functional assays. We found that some groups of candidate genes encoding putative subunits of the same phosphoprotein phosphatase complex also phenocopied each other, with RNAi knockdown resulting in statistically significant enhanced susceptibility to the same environmental stressor. This result was interpreted as evidence of a putative physical interaction between the gene products, and it is the basis for the identity of the subunits that we propose as constituents of the phosphoprotein phosphatase complexes that regulate DAF-16 during aging.

Our data support the possibility of multiple versions of the PP2A/4/6 complexes that are co-expressed during aging in worms that have specialized and partially overlapping functions in preserving health by conferring resistance to environmental insults (Fig. 7). Our results indicate that a PP2A complex comprised of the catalytic subunit LET-92, the scaffolding subunit PAA-1, and the regulatory subunit PPTR-1 is important for DAF-16-mediated resistance to ultraviolet irradiation in adult animals. In like manner a PP4 complex consisting of the catalytic subunit PPH-4.1/4.2, and the regulatory subunits SMK-1 and PPFR-2 appears to be necessary for DAF-16-mediated innate immunity later in life. The PP6 complex PPH-6/SAPS-1 also contributes to host defense in adult animals, but it appears to do so without affecting the transcriptional activity of DAF-16. The genetic and functional evidence that form the basis of our claims regarding these complexes is further substantiated by the fact that in each instance, the constituent subunits are known to be expressed together in the same tissues in *C. elegans*, and colocalization to the same subcellular compartment has been experimentally verified for several (Table S4). Different versions of the PP2/4/6 complexes may exist in the same cells simultaneously. Alternatively, the composition of the complexes may be dynamic such that individual regulatory subunits associate and dissociate according to cellular demands or in response to external stimuli. Future biochemical studies involving isolation of the complexes from wildtype postreproductive adult *C. elegans* and *in vitro* binding assays will validate the conclusions regarding the molecular composition of PP2A/4/6 complexes based on data from our genetic approach.

**Figure 7.**
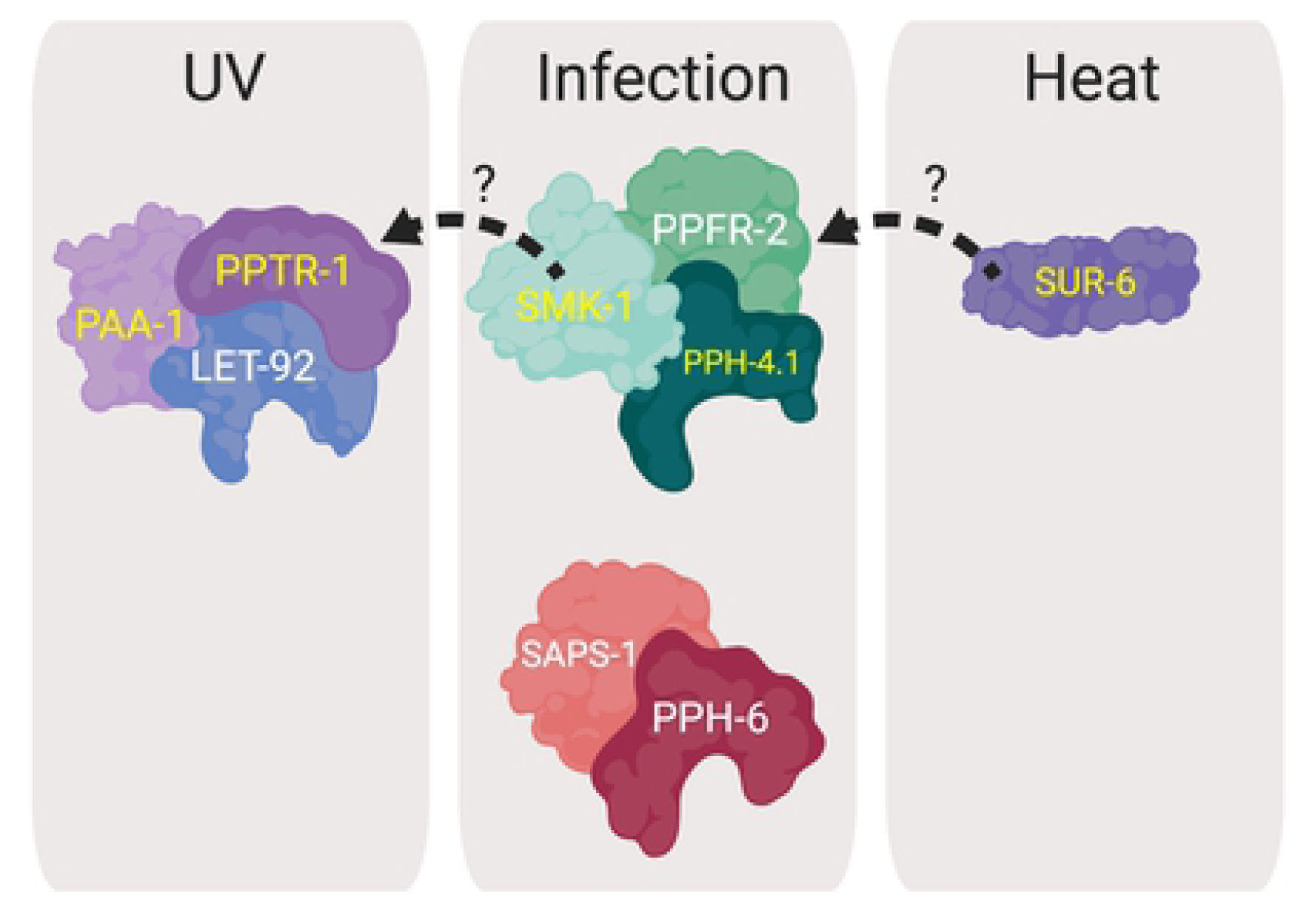
Proposed identity of PP2/4/6 complexes in *C. elegans* and their functions during aging. Our data support the existence of at least three central phosphoprotein phosphatase complexes during aging in *C. elegans*, but different combinations of subunits allow for more. The subunit composition of each complex of the PP2/4/6 family in adult *C. elegans* is indicated, according to the following color scheme: purple, PP2A subunits; green, PP4 subunits; red, PP6 subunits. Proteins whose names are labeled in yellow text are required for the age-dependent increase in DAF-16 transcriptional activity as measured in our *in vivo* reporter assay. While the PP2A complex is important for resistance to UV light, both the PP4 and PP6 complexes function in innate immunity in adult worms. Subunit exchange (indicated by dashed arrows) may take place between constituents of the PP2A and PP4 complexes such that SMK-1 associates with a version of the PP2A complex to confer resistance to UV irradiation and SUR-6 associates with PP4 to contribute to host defense. SUR-6 was the only PP2A/4/6 family member found to play a role in thermotolerance during adulthood. Protein structures depicted in this cartoon are for illustrative purposes only.

Our observations both confirm and extend previous reports of either physical or regulatory interactions between DAF-16 and the PP2A or PP4 complexes in *C. elegans* and in mammals. For example, PPTR-1 promotes the dephosphorylation of AKT-1 and is required for the extended lifespan and stress resistance phenotypes of *daf-2* mutants (21). These data strongly implicate PP2A as an indirect DAF-16 regulator in *C. elegans*, but no other PP2A complex members were isolated in the screen for *daf-2* suppressors that yielded *pptr-1*. Our data indicate that LET-92 and PAA-1 complete that tripartite PP2A complex that regulates DAF-16. Supporting this possibility, mammalian orthologues of the same three PP2A subunits that we identified were found to be binding partners of FoxO3a in HEK293 cells (109). Therefore, although the specific substrates may differ between *C. elegans* and mammals, regulation of FoxO transcription factors by PP2A appears to be evolutionarily conserved. The PP2A complex was not the first member of the PP2A/4/6 family to be identified as a regulator of DAF-16 activity. That distinction belongs to the PP4 complex and to SMK-1 in particular. Similar to the effect of inhibiting PPTR-1, loss-of-function mutations in *smk-1* abrogate *daf-2* phenotypes in larvae and young adults, including the elevated expression of DAF-16 transcriptional targets (43). More recently, RNAi inhibition of the PP4 catalytic subunits *pph-4.1/4.2* were also found to partially suppress *daf-2* (45). These subunits along with the regulatory subunit PPFR-2 copurify with SMK-1 from wild type, *daf-2* (e1370) and *daf-18* (mg198) animals, strongly suggesting that they form a complex *in vivo* (45). Our functional analysis provides an independent line of evidence to support this association between PPH-4.1/4.2, SMK-1 and PPFR-2.. Further, our data indicate that this particular version of the PP4 complex plays an important role during aging in wild type animals to stimulate the transcriptional activity of DAF-16.

Consistent with characterization of the PP4 catalytic subunits PPH-4.1 and PPH-*4.2* in *daf-2* mutants, we found that both are required to confer stress resistance and to activate DAF-16 transcriptional activity in adult animals. RNAi inhibition of either *pph-4.1* or *pph-4.2* resulted in two significant phenotypes in Day 6 adults: reduced expression of the *plys-7::GFP* reporter and enhanced susceptibility to *P. aeruginosa* infection. While knockdown of each gene had a similar effect on *plys-7::GFP* expression levels, RNAi targeting *pph-4.2* caused a more severe pathogen susceptibility phenotype. We also observed that knocking down *pph-4.2* but not *pph-4.1* resulted in a mild but statistically insignificant susceptibility to ultraviolet irradiation (Table 2; Fig S5). This may indicate that the two proteins have only partially overlapping functions. Taken together, our studies indicate that PPH-4.1 and its paralog PPH-4.2 are important during aging and are likely to be coexpressed in adult *C. elegans*.

Interestingly, in *C. elegans* the PP2A scaffold PAA-1 seems to associate with the PP4 subunit SMK-1 (45). Whether *paa-1* suppresses *daf-2* as does *smk-1* inhibition is unknown, yet in light of the biochemical evidence, one model is that PAA-1 could be included as a fourth subunit of the PP4 complex in *C. elegans*. Alternatively, there may be hybrid tripartite holoenzymes where two members are canonical subunits of one complex (e.g. PP2A) and the third member is considered to be a canonical subunit of a different complex (e.g. PP4). This would require certain regulatory or scaffolding subunits, perhaps including PAA-1, to have the ability to be incorporated into both PP2A and PP4 complexes. Regardless of their specific stoichiometry, biochemical evidence from mammalian systems supports the possibility of interchange of subunits between complexes of the PP2A/4/6 family. In affinity purification or immunoprecipitation experiments HEK293 cells, PP4c has been isolated as part of complexes with PP2A subunits, including orthologs of PAA-1, SUR-6, and PPTR-2 (51,52,110). Such heterotypic associations between members of the PP2A and PP4 complexes could explain two scenarios that we encountered in our studies where a singular representative of one PP complex shares a susceptibility phenotype with three members that are all part of a second PP complex. In particular, we found Day 6 adults to be susceptible to ultraviolet irradiation upon treatment with RNAi targeting not only the PP2A complex members *let-92*, *pptr-1*, and *paa-1* but also the PP4 subunit *smk-1*. Similarly, *sur-6* was the only PP2A subunit whose knockdown resulted in enhanced susceptibility to bacterial infection, as did RNAi targeting the PP4 complex members *pph-4.1/4.2*, *smk-1*, and *ppfr-2*. In each case a regulatory subunit seems to be operating separately from its cognate catalytic subunit. These results suggest that there may be alternative forms of the PP2A and PP4 complexes that include at least one regulatory subunit of the family that is not typically associated with that complex’s eponymous catalytic subunit.

In some cases we found that a gene’s requirement for the age-dependent increase in *plys-7::GFP* expression was apparently uncoupled from any role that it may have in conferring resistance to stress during adulthood. For example, neither *let-92* nor *ppfr-2* were required for the increase in *plys-7::GFP* expression in adult animals, but each conferred resistance to stress. Since LET-92 appears to mediate resistance to ultraviolet irradiation and PPFR-2 functions in host defense, in this case the functional data do not seem to support the possibility of a subunit exchange to yield a LET-92/PPFR-2 complex. While the requirement for LET-92 in conferring resistance to UV irradiation in adult worms without an accompanying effect on *plys-7::GFP* expression could suggest a DAF-16-independent function, all other PP2A subunit orthologs in our study whose knockdown caused adults to be more sensitive to stress were also necessary for the age-dependent increase in *plys-7::GFP* expression. Considering these results and in light of the aforementioned evidence from mammalian systems establishing a regulatory interaction between PP2A and FoxO transcription factors, we favor a model in which LET-92 is the catalytic subunit of one or more PP2A complexes that regulate the function of DAF-16 during aging, including its role in protecting animals from environmental insults. In another instance of incongruent phenotypes, knockdown of two putative PP2A B’’ regulatory subunits, F43B10.1 and T22D1.5, decreased expression levels of *plys-7::GFP* in Day 6 adults without enhancing the sensitivity of adult animals to environmental stress. Perhaps PP2A complexes that include these two subunits regulate DAF-16 in such a manner as to affect its transcriptional output. While *lys-7* would still be among the transcriptional targets of DAF-16 regulated by such a PP2A complex, other transcriptional targets including genes that are critical for mediating stress resistance would not be. Another possible explanation of this observation is that T22D1.5 and F43B10.1 may have redundant roles in directing DAF-16 to partially overlapping subsets of its transcriptional targets that include *lys-7*. A loss-of-function in either T22D1.5 or F43B10.1 alone would affect the transcriptional output of DAF-16 without completely abrogating DAF-16-mediated stress resistance. Only when the function of both regulators is diminished at the same time would DAF-16-mediated stress resistance be compromised because a critical combination of effectors fails to be upregulated. Testing this possibility would require further functional analyses of double mutants along with transcriptomic studies to determine how the transcriptional targets of DAF-16 during aging might change in the absence of PP2A complex members.

In the course of characterizing the entire PP2A/4/6 subfamily in our studies, we uncovered a new role for the PP6 complex in innate immunity during aging in *C. elegans*. Prior to our work on PPH-6, the *C. elegans* ortholog of PP6c, it had only been described in the context of development. In worm embryos PPH-6 is required for spindle positioning, similar to the role of its counterpart PP6c in mammals that controls spindle formation and the condensation, alignment, and segregation of chromosomes during mitosis (79,81,82,111). Based on the other functions of PP6c, we expected that PPH-6 might be necessary to protect adult *C. elegans* from ultraviolet irradiation. PP6c functions in both non-homologous end joining (NHEJ) and homology-directed repair of DNA, and without it cells are more sensitive to radiation (95, 100). We found no evidence to suggest that either PPH-6 or its regulatory subunit SAPS-1 confer resistance to UV in Day 6 animals. Instead, both PPH-6 and SAPS-1 were required for the survival of animals challenged with *P. aeruginosa*. Since the expression of immune effectors is the principle means of host defense in *C. elegans*, we speculate that PP6 regulates some aspect of that process either at an upstream step in immune signaling pathways or further downstream to influence the transcription of genes encoding antimicrobial products.

As the predominant antagonists to a slew of kinases, the catalytic activity of the PP2A/4/6 family is tightly controlled. Regulators of PP2A/4/6 may influence the composition of the holoenzyme, directly modify the active site of the catalytic subunit, or modulate the stability of the complexes. In some cases, the same regulatory protein can act at multiple levels. Since the substrate specificity of the PP2A/4/6 complexes is determined by the regulatory (B) subunits incorporated into the holoenzyme, one common mechanism of regulation involves modulating the assembly or the composition of the complexes. For example, some regulators of the PP2A/4/6 family such as CIP2A or Greatwall directly bind to the B or the A (scaffolding) subunits to prevent their incorporation into the multiprotein complexes (77, 112). Other regulators add or remove posttranslational modifications to the catalytic subunit that influence its affinity for particular regulatory subunits. While phosphorylation tends to disrupt the interaction between the catalytic and regulatory subunits, methylation enhances the binding of the catalytic subunit to B and B’ subunits (113, 114). The catalytic subunit is susceptible to additional regulation specifically at the active site. In particular, during its biogenesis, PP2Ac is bound by PTPA which helps to configure the active site for serine/threonine phosphatase activity (53, 115). Later, PP2Ac and PP4c may be inactivated by PME1 or by the coordinated function of TIPRL and α4 (homologues of *S. cerevisiae* Tip41p and Tap42p, respectively) that expel metal ions from the active site (58,59,116). When bound by these negative regulators, catalytic subunits or holoenzymes are maintained in a stable latent state. Our studies of the *C. elegans* orthologs of TIPRL (ZK688.9) and α4 (*ppfr-4*) during aging were inconclusive since RNAi treatments targeting them failed to produce statistically significant phenotypes in our stress assays. We did, however, observe a nearly significant increase in the sensitivity of Day 6 adults to UV radiation when *ppfr-4* was knocked down. Taken together with our other data, this could indicate that PPFR-4 plays a role in activating the LET-92/PAA-1/PPTR-1 PP2A complex during aging in *C. elegans*. Indeed, there is some evidence to suggest that α4 is capable of increasing the activity of PP2Ac in mammals (117, 118). This may be at least partly attributable to the function of α4 in protecting PP2Ac from ubiquitin-mediated proteasomal degradation (119, 120). Since knocking down ZK688.9 did not recapitulate phenotypes associated with RNAi inhibition of *ppfr-4* in our hands, we are unable to say whether their gene products are likely to cooperate as they do in mammals. One possibility, though, is that the interplay between these proteins in *C. elegans* is more similar to the situation in *Saccharomyces cerevisiae* where Tip41p and Tap42p function independently and antagonistically (121). Further experiments will be necessary to investigate the regulatory mechanism governing the activity of the PP2A/4/6 family during aging in worms and other species.

Our work establishes roles for the PP2A, PP4, and PP6 phosphatases in contributing to the healthspan of postreproductive adult *C. elegans* through conferring resistance to environmental insults at least in part by modulating transcription. The mechanistic basis for this function will be revealed when the substrates of the complexes in aging animals are uncovered. In light of evidence from mammalian systems, during aging in *C. elegans* the PP2A and PP4 complexes may act to dephosphorylate DAF-16 itself or its upstream inhibitory kinases. Yet if this were true and the phosphatases regulate an early step in DAF-16 activation to counteract inhibition through IIS, we may have expected to find significant functional overlap between the PP2A and PP4 complexes such that both would have conferred resistance to the same stresses. Instead, we found that each complex seems to specialize in contributing to the resistance to particular insults. This suggests that PP2A and PP4 may regulate DAF-16 at a more downstream step, possibly functioning to tailor the transcriptional output of DAF-16 to include only discrete subsets of its complete repertoire of targets. Consistent with this possibility, the transcription elongation factor SPT-5 was recently identified as a substrate of PP4 in *daf-2 (e1370)* mutants where it plays a role in recruiting RNA PolII to the promoters of some but not all of the genes regulated by DAF-16 (45). Should the PP2A/4/6 family act at a similar level in adult wildtype animals, there are other means by which its members are known to influence gene expression that may be of particular relevance to aging. Across evolutionarily diverse species, these protein phosphatases regulate histone modification, chromatin organization, and mRNA processing which all undergo significant changes over time (63,64,89,106,122–125). Since we and others find that the PP2A/4/6 phosphatases are important for promoting healthspan, it may be by acting through these pathways that the PP2A/4/6 family modulates the changes in gene expression that are necessary to uphold vitality.

## Acknowledgments

We thank Jennifer Goldfarb for assistance with data collection. Kali Carrasco and Patrick Mitrano-Towers provided valuable feedback during the preparation of the manuscript. We are grateful to members of the staff of the Villanova Department of Biology for their technical assistance.

## Supporting Information

**Figure S1. The catalytic subunits of the PP2A/4/6 subfamily of phosphoprotein phosphatases are highly conserved between *C. elegans* and humans.** The amino acid sequences of *C. elegans* LET-92, PPH-4.1, PPH-4.2 isoform a, and PPH-6 were aligned to their human orthologs PP2Ac, PP4c, and PP6c using Clustal Omega. Residue numbers are indicated on the right. The degree of conservation of individual amino acids across all seven proteins is denoted by symbols where asterisks (*) indicate identical residues, two dots (:) indicate highly similar residues, and one dot (.) indicates somewhat similar residues at a particular position. Characteristic domains of the subfamily are indicated, including the helix switch, loop switch, and TPDYFL motif (31). Specific residues associated with metal ion coordination and catalysis as well as those that interact with regulatory proteins are denoted by colored symbols according to the legend (115,116,126,127). In cases where multiple annotations apply to the same residue, corresponding symbols are vertically stacked and squares may be split diagonally.

**Figure S2. Interaction network for putative subunits of the *C. elegans* PP4 complex reveals possible connections to other members of the PP2A/4/6 subfamily.** A list of potential subunits of the *C. elegans* PP4 complex including PPFR-1, PPFR-2, PPH-4.1, and SMK-1 in addition to the regulatory protein PPFR-4 was submitted for analysis to Genemania (genemania.org). The resulting network diagram depicts multiple types of interactions including physical interactions (red lines), predicted interactions (orange lines), or genetic interactions (green lines). Other similarities between proteins including co-expression (purple lines) or shared protein domains (yellow lines) are also indicated. In cases where only a Wormbase gene identifier or a CDS identifier was listed in the network diagram, the *C. elegans* gene name and/or the human ortholog (in parentheses) is provided in an adjacent grey box. Colored outlines surrounding nodes indicate that a particular protein is either an ortholog of a human PP2A/4/6 complex subunit or is an ortholog of a protein that regulates the activity of one or more members of the PP2A/4/6 family. Since including F46C5.6 as part of the query resulted in a second node that was not part of the larger network it was eliminated from the analysis.

**Figure S3. PP4 subunit homologues PPFR-1 and F46C5.6 do not function in innate immunity in *C. elegans*.** RNAi treatment targeting *C. elegans* homologs of regulatory subunits of the PP4 complex was initiated at the L1 stage. After knockdown of F46C5.6 (A,C) or *ppfr-1* (B,D), worms were infected at the L4 larval stage (A, B) or at Day 6 of adulthood (C, D). A representative plot of the fraction of worms alive at each time point after the infection was initiated is shown. In all cases RNAi targeting *daf-16* or *smk-1* and the empty RNAi vector L4440 were included as controls. Statistical analyses indicate that neither of the RNAi treatments had a significant effect on the survival of worms following bacterial infection.

**Figure S4. *C. elegans* homologues of human PP2A complex subunits with no apparent role in innate immunity during aging.** Representative survival curves for worms treated with RNAi to knockdown the indicated gene beginning at L1 and then infected with *P. aeruginosa* at the L4 larval stage (A-F) or at Day 6 of adulthood (G-N) are shown. Since RNAi inhibition of *let-92* and *paa-1* arrested larval development, knockdown of those genes was initiated at the L4 stage and worms were infected only at Day 6 (M,N). A representative plot of the fraction of worms alive at each time point after the infection began is shown. All plots include data for animals treated with the empty RNAi vector L4440 and for RNAi knockdown of *daf-16* and *smk-1*. Statistical analyses indicate that none of the RNAi treatments had a significant effect on the survival of worms following bacterial infection.

**Figure S5. Three regulatory subunits of the PP4 complex do not appear to function in conferring resistance to UV irradiation.** RNAi treatment targeting *C. elegans* homologs of regulatory subunits of the PP4 complex F46C5.6 (A, D), *ppfr-1* (B, E), and *ppfr-2* (C, F) was initiated at the L1 stage and continued for the duration of the assay. Worms were exposed to UV irradiation at the L4 larval stage (A-C) or at D6 of adulthood (D-F) after which their survival under standard culturing conditions was monitored. A representative plot of the fraction of worms alive at each time point following exposure to UV radiation is shown. All plots include data for animals treated with the empty RNAi vector L4440 and for RNAi knockdown of *daf-16* and *smk-1*. Statistical analyses indicate that none of the RNAi treatments had a significant effect on the survival of worms following UV irradiation.

**Figure S6. *C. elegans* homologues of human PP2A complex subunits with no apparent role in conferring resistance to UV irradiation.** Representative survival curves for worms treated with RNAi to knockdown the indicated gene beginning at L1 and irradiated with ultraviolet light at the L4 larval stage (A-F) or at Day 6 of adulthood (G-L) are shown. Following irradiation, worms were returned to standard culture conditions and their survival was monitored over time. A representative plot of the fraction of worms alive at each time point following exposure to UV radiation is shown. All plots include data for animals treated with the empty RNAi vector L4440 and for RNAi knockdown of *daf-16* and *smk-1*. Statistical analyses indicate that none of the RNAi treatments had a significant effect on the survival of worms following UV irradiation.

**Figure S7. The PP6 regulatory subunit SAPS-1 plays no role in protecting *C. elegans* from UV irradiation.** RNAi treatment targeting *C. elegans* homolog of the PP6 regulatory subunit *saps-1* was initiated at the L1 stage and continued for the duration of the assay. Worms were exposed to UV irradiation at the L4 larval stage (A) or at D6 of adulthood (B) after which their survival under standard culturing conditions was monitored. A representative plot of the fraction of worms alive at each time point following exposure to UV radiation is shown. All plots include data for animals treated with the empty RNAi vector L4440 and for RNAi knockdown of *daf-16* and *smk-1*. Statistical analyses indicate that RNAi targeting *saps-1* had no significant effect on the survival of worms following UV irradiation.

**Figure S8. PP4 subunit homologues PPFR-1 and PPFR-2 do not confer thermotolerance to *C. elegans*.** From L1 until death *C. elegans* were treated with RNAi targeting putative regulatory subunits of the PP4 complex *ppfr-1* (A, C) or *ppfr-2* (B,D). Worms were shifted from 20° C to 35° C at larval stage L4 (A, B) or D6 (C, D) and maintained at the high temperature until death. A representative plot of the fraction of worms alive at each time point during the incubation at 35° C is shown. In all cases RNAi targeting *daf-16* or *smk-1* and the empty RNAi vector L4440 were included as controls. Statistical analyses indicate that neither of the RNAi treatments had a significant effect on the survival of worms under heat stress.

**Figure S9. *C. elegans* homologues of human PP2A complex subunits with no apparent role in thermotolerance during adulthood.** From L1 until death worms were treated with RNAi targeting *C. elegans* orthologs of catalytic and regulatory subunits of the human PP2A complex. At either the L4 stage (A-F) or at Day 6 of adulthood (G-N) worms were shifted from the standard incubation temperature of 20° C to 35° C and were maintained at the high temperature until death. Since RNAi inhibition of *let-92* and *paa-1* arrested larval development, knockdown of those genes was initiated at the L4 stage and worms were shifted to high temperature only at Day 6 (M,N). A representative plot of the fraction of worms alive at each time point during the incubation at 35° C is shown. In all cases RNAi targeting *daf-16* or *smk-1* and the empty RNAi vector L4440 were included as controls. Statistical analyses indicate that none of the RNAi treatments had a significant effect on the survival of worms under heat stress.

**Figure S10. The PP6 regulatory subunit SAPS-1 plays no role in the response to heat stress.** From L1 until death *C. elegans* were treated with RNAi targeting *saps-1*, a subunit of the PP6 complex in *C. elegans*. Worms were shifted from 20° C to 35° C at larval stage L4 (A) or D6 (B) and maintained at the high temperature until death. A representative plot of the fraction of worms alive at each time point during the incubation at 35° C is shown. In all cases RNAi targeting *daf-16* or *smk-1* and the empty RNAi vector L4440 were included as controls. Statistical analyses indicate that RNAi targeting *saps-1* did not have a significant effect on the survival of worms under heat stress.

**Figure S11. *C. elegans* orthologs of regulators of the PP2/4/6 family do not contribute to host defense during adulthood.** Beginning at the L1 stage the *C. elegans* homologs of human PTPA (Y71H2AM.20; A, C) and TIPRL (ZK688.9; B, D) were targeted with RNAi. Worms were infected with *P. aeruginosa* at the L4 (A, B) larval stage or at D6 (C, D) of adulthood. A representative plot of the fraction of worms alive at each time point after the infection was initiated is shown. In all cases RNAi targeting *daf-16* or *smk-1* and the empty RNAi vector L4440 were included as controls. Statistical analyses indicate that none of the RNAi treatments had a significant effect on the survival of worms following bacterial infection.

**Figure S12. *C. elegans* orthologs of PTPA and TIPRL have no apparent function in conferring resistance to UV irradiation.** RNAi treatment targeting the PTPA ortholog Y71H2AM.20 (A, B) or the TIPRL ortholog ZK688.9 (C, D) was initiated at the L1 stage and continued for the duration of the assay. Worms were exposed to UV irradiation at the L4 larval stage (A, C) or at D6 of adulthood (B, D) after which their survival under standard culturing conditions was monitored. A representative plot of the fraction of worms alive at each time point following exposure to UV radiation is shown. All plots include data for animals treated with the empty RNAi vector L4440 and for RNAi knockdown of *daf-16* and *smk-1*. Statistical analyses indicate that RNAi targeting Y71H2AM.20 or ZK688.9 had no significant effect on the survival of worms following UV irradiation.

**Figure S13. *C. elegans* orthologs of regulators of the PP2/4/6 family do not affect susceptibility to heat stress.** Beginning at the L1 stage the *C. elegans* homologs of human α4 (*ppfr-4*; A, D), PTPA (Y71H2AM.20; B, E), and TIPRL (ZK688.9; C, F) were targeted with RNAi. Worms were shifted from 20° C to 35° C at larval stage L4 (A-C) or D6 (D-F) and maintained at the high temperature until death. A representative plot of the fraction of worms alive at each time point during the incubation at 35° C is shown. In all cases RNAi targeting *daf-16* or *smk-1* and the empty RNAi vector L4440 were included as controls. Statistical analyses indicate that none of the RNAi treatments had a significant effect on the survival of worms under heat stress.

**Supplemental Table 1. Percent identity matrix for *C. elegans* and human PP2A/4/6 catalytic subunits.** Output from Clustal Omega alignment of *C. elegans* and human catalytic subunits of the PP2A/4/6 subfamily showing percent identity between each pairwise comparison. The best matching ortholog in each column (highest percent identity) is highlighted in green.

**Supplemental Table 2. PPH-4.1 interactors.** Predicted and experimentally validated interactors of PPH-4.1 as listed in the Wormbase protein interaction database are shown.

**Supplemental Table 3. SMK-1 interactors.** Predicted and experimentally validated interactors of SMK-1 as listed in the Wormbase protein interaction database are shown.

**Supplemental Table 4. Location and timing of expression of *C. elegans* PP2A/4/6 genes.** Genes listed in this table correspond to the following three groups: 1) those encoding members of the holoenzyme complexes that we propose in Figure 7; 2) T22D1.5 and F43B10.1, orthologs of PP2B” regulatory subunits that we found to be necessary for the age-dependent increase in DAF-16 transcriptional activity; and 3) orthologs of regulatory proteins. *daf-16* expression information is included as a reference. Expression of genes in tissues where DAF-16 function has been explicitly tested in previous studies, including body wall muscle, intestine, and neurons, are highlighted by coloring the names of those tissues (red, green, and blue, respectively) in the “Anatomical sites of expression” column.

